# *Os*r40g3 imparts salt tolerance by regulating GF14e-mediated gibberellin metabolism to activate EG45 in rice

**DOI:** 10.1101/2020.04.26.062158

**Authors:** Chandan Roy, Salman Sahid, Dibyendu Shee, Riddhi Datta, Soumitra Paul

## Abstract

Under changing environmental conditions, salt stress has posed a severe threat to agriculture. Although the R40 family lectins are known to be associated with osmotic stress response, their mechanism of action remains elusive. Among them, *Osr40g3* displays the highest expression under salt stress. Here, we report that the constitutive overexpression of *Osr40g3* imparts salt tolerance but displays pollen sterility and poor seed development in rice. Promoter analysis and gene expression studies revealed that the gene follows a precise tissue-specific expression pattern, which is essential for proper seed development. Overexpressing the gene under the control of its native promoter rescued the pollen-sterile phenotype while significantly improving salt tolerance. Protein-protein interaction studies demonstrated that *Os*r40g3 positively regulates an expansin protein, *Os*EG45, while decreasing the stability of a 14-3-3 protein, *Os*GF14e. Correspondingly, the *OsEG45* overexpression and *OsGF14e* silencing lines display a salt-tolerant phenotype. Again, silencing *OsEG45* in the background of *OsGF14e* silencing lines resulted in a salt-sensitive phenotype, indicating that salt tolerance of the *OsGF14e* silencing lines is *OsEG45*-dependent. Notably, the *OsGF14e* gene displays early salt responsiveness, while *Osr40g3* and *OsEG45* display a late response, indicating a spatio-temporal regulation of these genes. Interestingly, constitutive overexpression of *Osr40g3* or silencing of *OsGF14e* leads to diminished gibberellic acid (GA) accumulation that activates the *OsEG45* gene. Together, our study demonstrates that during salt stress, *Osr40g3*, a late salt-responsive gene, confers salt tolerance by negatively regulating *Os*GF14e while positively regulating *Os*EG45 via a GA-mediated pathway. This mechanistic insight broadens our understanding of lectin-mediated regulation of salt tolerance.

## INTRODUCTION

Plants are sessile organisms that encounter environmental challenges *in situ*. Fascinatingly they can counteract most of these threats with their intricate defense signaling network. Lectins are carbohydrate-binding proteins that are excellent candidates in this signaling network and modulate biotic and abiotic stress responses in different plant species (Lannoo and van Damme, 2014; Sahid *et al*., 2020).

With the changing environmental conditions, salt stress has become a serious threat to crop productivity. The effect of salt stress in rice growth is primarily associated with various morphological alterations like rolling of leaves, delay of panicle emergence and anthesis, spikelet desiccation, loss of pollen viability, etc. Since hyper-salinity reduces soil water potential, it hampers water absorption in plants. To survive under this condition, plants accumulate different osmoprotectants, compatible solutes, polyamines, and secondary metabolites, regulate ion homeostasis, and activate antioxidant systems (Golldack *et al*., 2014). An intricate phytohormone signaling network modulates these metabolic changes. The role of abscisic acid (ABA) in regulating salt stress is well documented (Verma *et al*., 2016). However, emerging evidence supports the role of other plant growth regulators, like gibberellic acid (GA), in relieving the effect of osmotic stress (Achard *et al*., 2006; Verma *et al*., 2016). Intriguingly, the lower accumulation of GA is associated with higher salt tolerance and reduced plant growth. For example, GA deficiency imparts drought tolerance with reduced plant height and lodging in teff and finger millet (Plaza-Wüthrich *et al*., 2016). The *SPINDLY* gene, encoding a negative regulator of GA, is induced in response to osmotic stress in Arabidopsis (Qin *et al*., 2011). On the other hand, the quadruple DELLA mutant lacking *GAI, RGA, RGL1*, and *RGL2* exhibits less growth inhibition in Arabidopsis under salt stress suggesting a DELLA-dependent growth restrain under salt stress that is also associated with lowering of the bioactive GA levels (Verma *et al*., 2016).

Among various stress-responsive proteins, lectins are gaining importance due to their diverse regulatory role against environmental challenges (Lannoo and van Damme, 2014). Plant lectins can be classified into 12 protein families (Jiang *et al*., 2010). Among them, the R40 family plays a crucial role in regulating drought and salt stress (Moons *et al*., 1995; 1997). In rice, the family consists of five functional members, namely *Os*r40c1, *Os*r40c2, *Os*r40g2, *Os*r40g3, and putative *Os*r40c1. Apart from osmotic stress, ABA, methyl jasmonate, and ethephone treatments can also induce the expression of these genes (Moons *et al*., 1995; 1997). Recently, we demonstrated that overexpression of *Os*r40c1 can impart drought tolerance in rice (Sahid *et al*., 2020). However, the mechanism of osmotic stress tolerance by the other R40 family proteins in rice remains elusive.

In this study, we aim to decipher the mechanism of how *Os*r40g3 regulates salt tolerance in rice. We observed that *Os*r40g3 exhibits dual function in positively regulating the expansin protein *Os*EG45 while negatively regulating the 14-3-3 family protein *Os*GF14e. The diminished accumulation of *Os*GF14e lowers the GA level, further inducing *OsEG45* expression, thus leading to salt tolerance in rice.

## RESULTS

### Expression of *Osr40g3* positively correlated with the salt tolerance potential in rice

To investigate the salt tolerance potential of selected eight *indica* rice cultivars, we exposed the seedlings to salt stress for 5 d, and several morphological and biochemical parameters were recorded (Figure S1A, S2). Among them, Nonabokhra was the most tolerant cultivar, while MTU1010 was the most sensitive one. Interestingly, we observed that among all the five *OsR40* genes, the expression of *Osr40g3* was highest under salt stress (Figure S3). This clue prompted us to explore if *Osr40g3* expression correlates with the salt tolerance potential in rice. Indeed, we observed that expression of this gene under salt stress positively correlates with the salt tolerance potential of the eight rice cultivars with a 4.429-fold induction in the tolerant Nonabokhra cultivar against a 2.218-fold induction in the sensitive MTU1010 cultivar (Figure S1B).

### Overexpression of *Osr40g3* improved salt tolerance but impaired seed development in rice

To validate the function of *Osr40g3* in salt tolerance, we generated *Osr40g3* overexpressing rice plants (*ox_c_* lines) harboring the recombinant *35S::Osr40g3* construct. In the vegetative stage, the plants displayed no phenotypic abnormalities except a reduced plant height. Surprisingly, however, the lines developed a shorter panicle length and failed to produce seeds (Figure 1A-L). We carried out plant transformation in four independent batches. Only two transgenic plants (out of 13) from the first batch and three (out of 11 and 15, respectively) from each of the second and third batches produced 30-35 seeds per plant. Twenty six independent transgenic lines were raised in the fourth batch and were used for stress assay.

**Figure 1.**
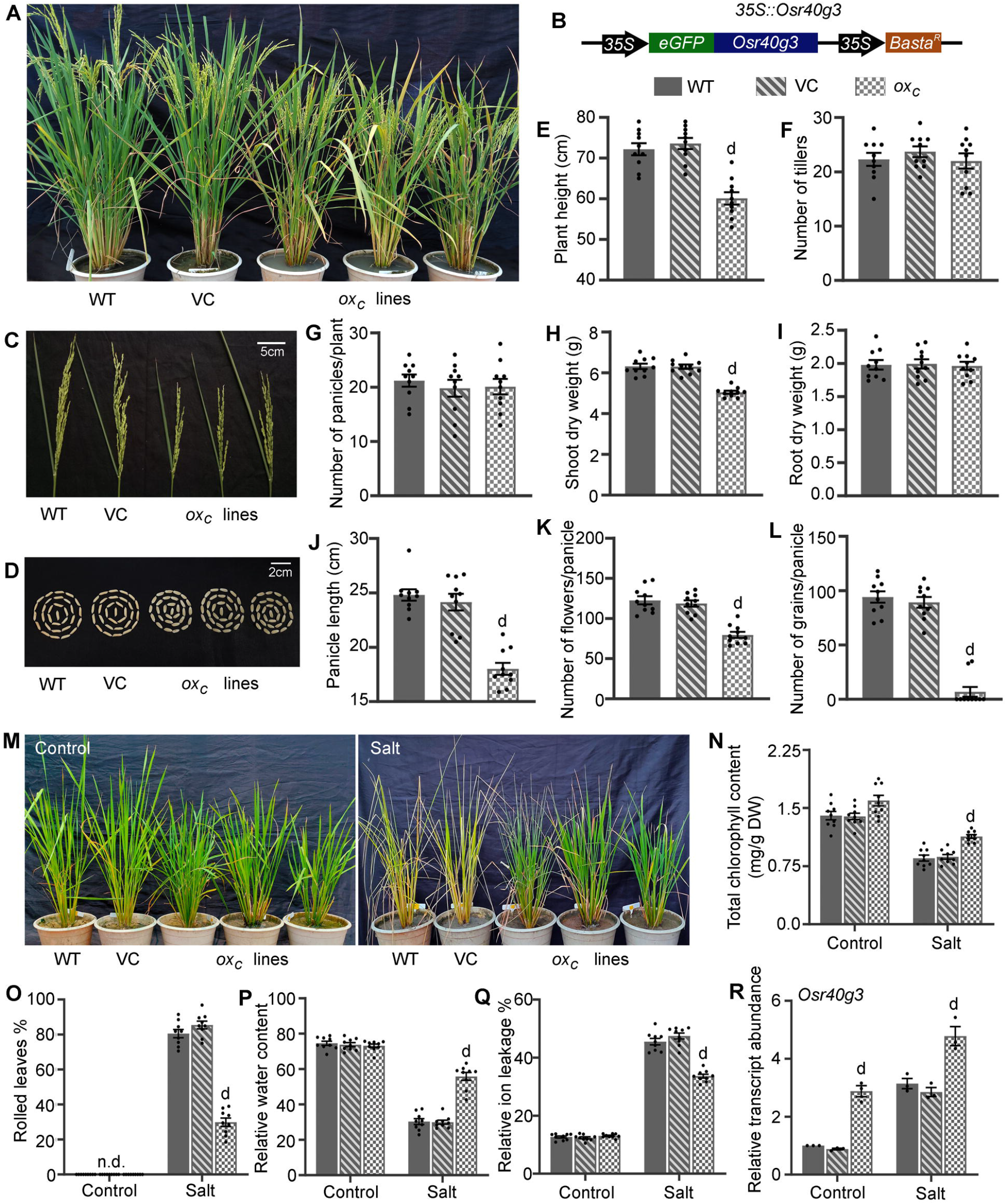
Transgenic rice lines constitutively overexpressing *Osr40g3* (*ox_c_*) displayed improved salt tolerance but impaired seed development. Different morphological parameters of the transgenic lines harboring the *35S::Osr40g3* construct (*ox_c_* lines) along with WT and VC (*35S::GFP*) plants were analyzed. The WT, VC and transgenic *ox_c_* lines were exposed to salt stress for 5 d and analyzed. (A) WT, VC and 3 representative independent *ox_c_* lines at reproductive stage, (B) overexpression construct, (C) panicles, (D) grain morphology (empty grains in transgenic lines), (E) plant height, (F) number of tillers, (G) numbers of panicles/plant, (H) shoot dry weight, (I) root dry weight, (J) panicle length, (K) numbers of flowers/panicle, (L) number of grains/panicle, (M) morphological response of WT, VC and 3 representative independent *ox_c_* lines under salt stress, (N) total chlorophyll content, (O) rolled leaf percentage, (P) relative water content, (Q) relative ion leakage percentage, and (R) relative transcript abundance of *Osr40g3* gene in response to salt stress. Ten independent transgenic lines (T_0_) were considered for morphological analysis. The stress assay was independently repeated thrice with 3 independent *ox_c_* lines (T_0_) in each set. Results were represented as mean ± SEM and the statistical difference between WT, VC and transgenic lines was denoted by ‘d’ at P<0.0001. n.d.: not detected.

On exposure to salt stress, the transgenic plants displayed significantly improved tolerance over the wild-type (WT) and vector control (VC) plants (Figure 1M). The transgenic plants maintained lower rolled leaf percentage, ion leakage, and chlorophyll degradation but higher relative water content (RWC) under salt stress (Figure 1N-Q). They accumulated significantly elevated levels of osmolytes but displayed lower lipid peroxidation and H_2_O_2_ accumulation (Figure S4A-D). We observed that the expression of the *Osr40g3* gene was significantly higher in the transgenic lines under control as well as salt stress which could account for their tolerant phenotype (Figure 1R). The amount of sodium (Na^+^), potassium (K^+^), and chloride (Cl^-^) ions as well as electrical conductivity (EC) of the soil indicated its saline nature after salt treatment (Table S1). Lower accumulation of Na^+^ and higher K^+^ level in the transgenic plants during salt stress also indicated their tolerant phenotype (Figure S4E-F).

Surprisingly, the transgenic *Arabidopsis* plants (*ec* lines) ectopically expressing the *Osr40g3* gene did not display any phenotypic abnormality (Figure S5). Yet, similar to rice, they exhibited significantly improved salt tolerance (Figure S6). These observations confirmed the role of *Osr40g3* in imparting salt tolerance in plants but indicated that its interference with seed development is rice-specific.

### Constitutive overexpression of *Osr40g3* resulted in pollen sterility in rice

To understand the differential interference of *Osr40g3* in seed development, we analyzed the floral morphology of both the rice and *Arabidopsis* transgenic plants. In rice, flowers of *ox_c_* plants were shorter than WT and VC flowers (Figure 2A-C). Further, the pollen viability in all *ox_c_* flowers was drastically reduced (Figure 2D-E). *In vitro* pollen germination assay displayed that most of the pollen grains from *ox_c_* plants failed to produce pollen tubes (Figure 2F). On the contrary, the transgenic *Arabidopsis* plants showed no alteration in floral morphology, pollen viability, and pollen germination (Figure S7).

**Figure 2.**
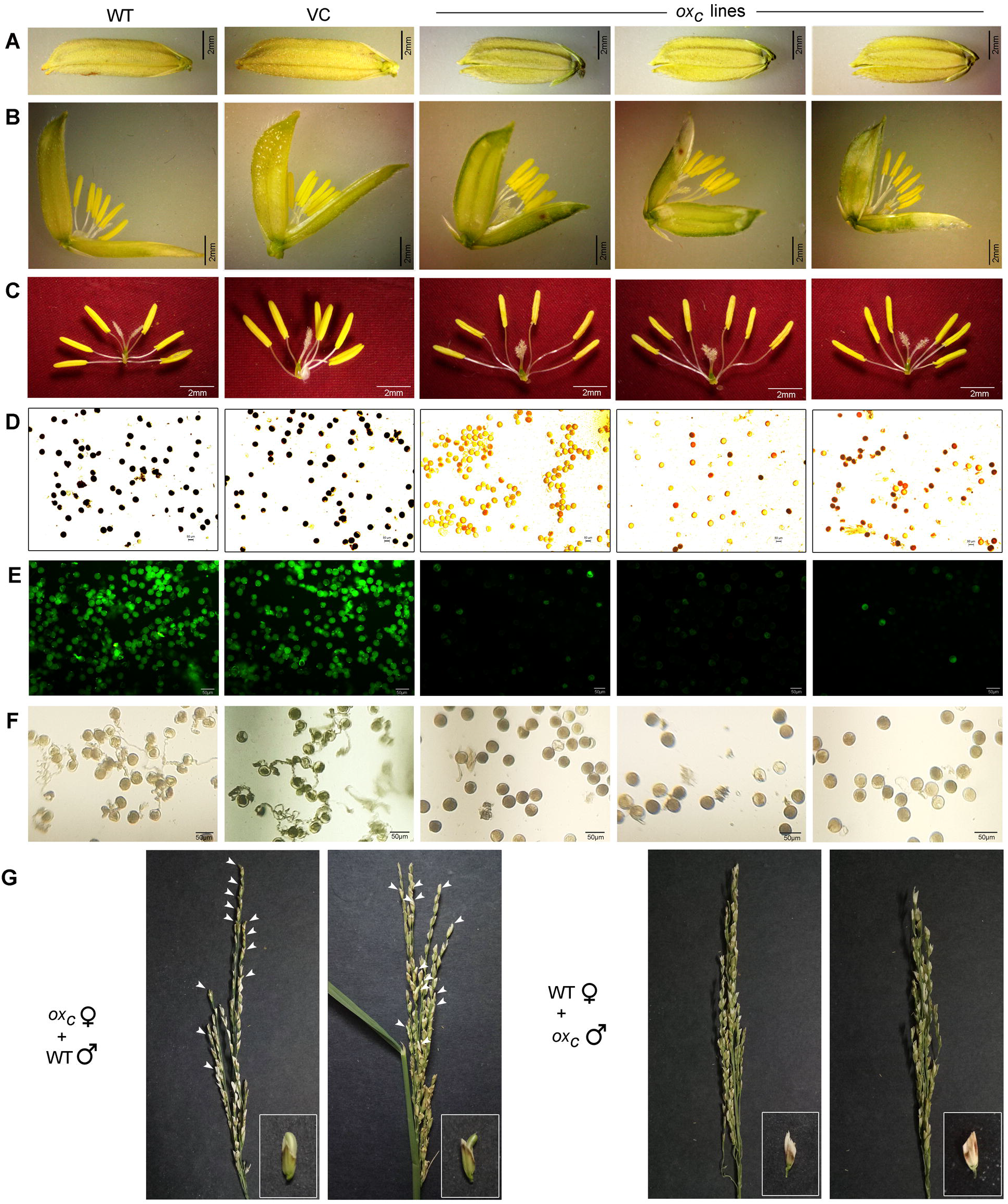
Floral morphology of transgenic rice lines constitutively overexpressing *Osr40g3* (*ox_c_*) and hybridization studies. The floral morphology and pollen viability of 3 representative independent transgenic lines harboring the *35S::Osr40g3* construct (*ox_c_* lines) along with the WT and VC (*35S::GFP*) plants were analyzed. (A) Entire flower, (B) dissected flower, (C) stamens with pistil, (D) pollen viability by I_2_-KI staining method, (E) pollen viability by FDA staining method, and (F) *in vitro* pollen germination assay. For floral morphology, 3 flowers from each of 10 independent T_0_ transgenic lines were considered. For pollen viability and *in vitro* germination assay, at least 500 pollen grains from each line were studied. (G) Crossing of emasculated transgenic *ox_c_* plants with WT pollen grains and crossing of emasculated WT plants with the pollens from transgenic *ox_c_* plants. The experiment was independently repeated in three sets. Arrows indicate hybrid seeds.

To check if the failure to produce seeds in the transgenic rice lines was due to pollen sterility alone, we crossed the *ox_c_* plants with WT plants in two separate hybridization programs. When the *ox_c_* plants were used as the male parents and hybridized with the emasculated WT plants, no seeds developed, confirming the pollen sterility in transgenic plants. Interestingly, when *ox_c_* plants were used as the female parent and crossed with WT plants as the male parent, hybrid seeds were developed, thus, indicating no defect in the pistil (Figure 2G).

### *Osr40g3* displayed a strict tissue-specific expression pattern in rice

To understand if the *Osr40g3* gene has any specific expression pattern in rice, we analyzed its expression in the WT plant from different tissues at different developmental stages. For this purpose, we used germinating seedlings, 60 d, and 120 d old plants and collected samples from the root, leaf, flag leaf, flower, stamen, pollen grains, and seeds, as applicable. We collected stamens from different stages of anthesis and seeds from the milk, dough, and mature stages. The *Osr40g3* gene displayed the highest expression level in leaves and no expression in stamens (Figure 3A). For further validation, we analyzed its promoter activity by histochemical GUS assay. GUS expression was observed in roots, leaves, pistils, and seeds but not in the stamens and pollen grains, supporting its tissue-specific expression pattern (Figure 3B-C). Further, we observed that the promoter activity of *Osr40g3* was strongly inducible under salt stress, thus suggesting its salt-responsive nature (Figure S8).

**Figure 3.**
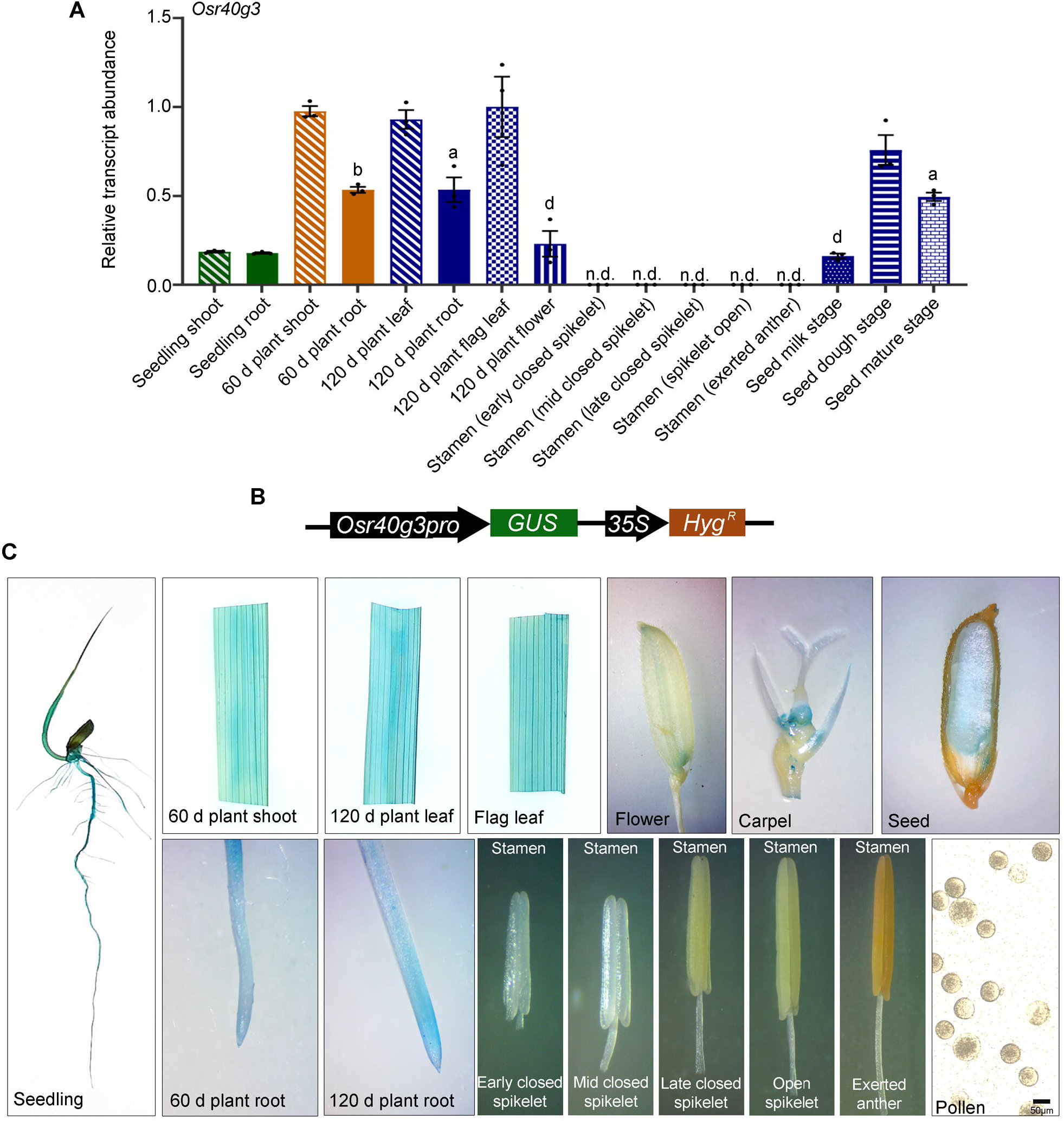
*Osr40g3* follows a precise tissue-specific expression pattern in rice. (A) Relative transcript abundance of *Osr40g3* gene in different tissues of WT rice plants was analyzed. The experiment was independently repeated thrice and results were represented as mean ± SEM. Statistical difference between different samples was denoted by different letters at P<0.05 (a), P<0.01 (b), and P<0.0001 (d). (B) The *Osr40g3pro::GUS* construct. (C) Representative image of histochemical GUS staining showing *Osr40g3* promoter activity in different tissues under various developmental stages. The experiment was independently repeated thrice. n.d.: not detected.

### Overexpression of *Osr40g3* under the control of native promoter imparted salt tolerance and rescued pollen sterility

Since *Osr40g3* followed a strict tissue-specific expression pattern, we generated transgenic rice plants (*ox_n_* lines) overexpressing the *Osr40g3* gene under the control of the native promoter. Out of the 14 independent transgenic lines, we selected three lines for further analysis. Unlike the *ox_c_* plants, these *ox_n_* plants displayed no phenotypic abnormalities (Figure 4A-C, S9). The *ox_n_* pollen grains also exhibited similar viability and pollen germination efficiency to the WT and VC plants (Figure S10). On the other hand, these *ox_n_* lines also displayed significantly improved salt tolerance and higher *Osr40g3* transcript abundance over the WT and VC plants (Figure 4D-I, S11). Next, we compared the salt tolerance potential of 30-day-old *ox_c_* and *ox_n_* lines in response to 5 d of salt exposure. Both *ox_c_* and *ox_n_* lines displayed improved salt tolerance with 100 % survival compared to 33.33 % in WT and 25 % in VC plants. However, the percentage of the rolled leaves was higher in the *ox_c_* lines compared to the *ox_n_* lines under salt stress (Figure 4J-K). To check if *Osr40g3* overexpression can alter expression of the other *OsR40* genes, we analyzed the relative transcript abundance in the *ox_c_* and *ox_n_* lines. However, we did not observe any differential expression (Figure S12).

**Figure 4.**
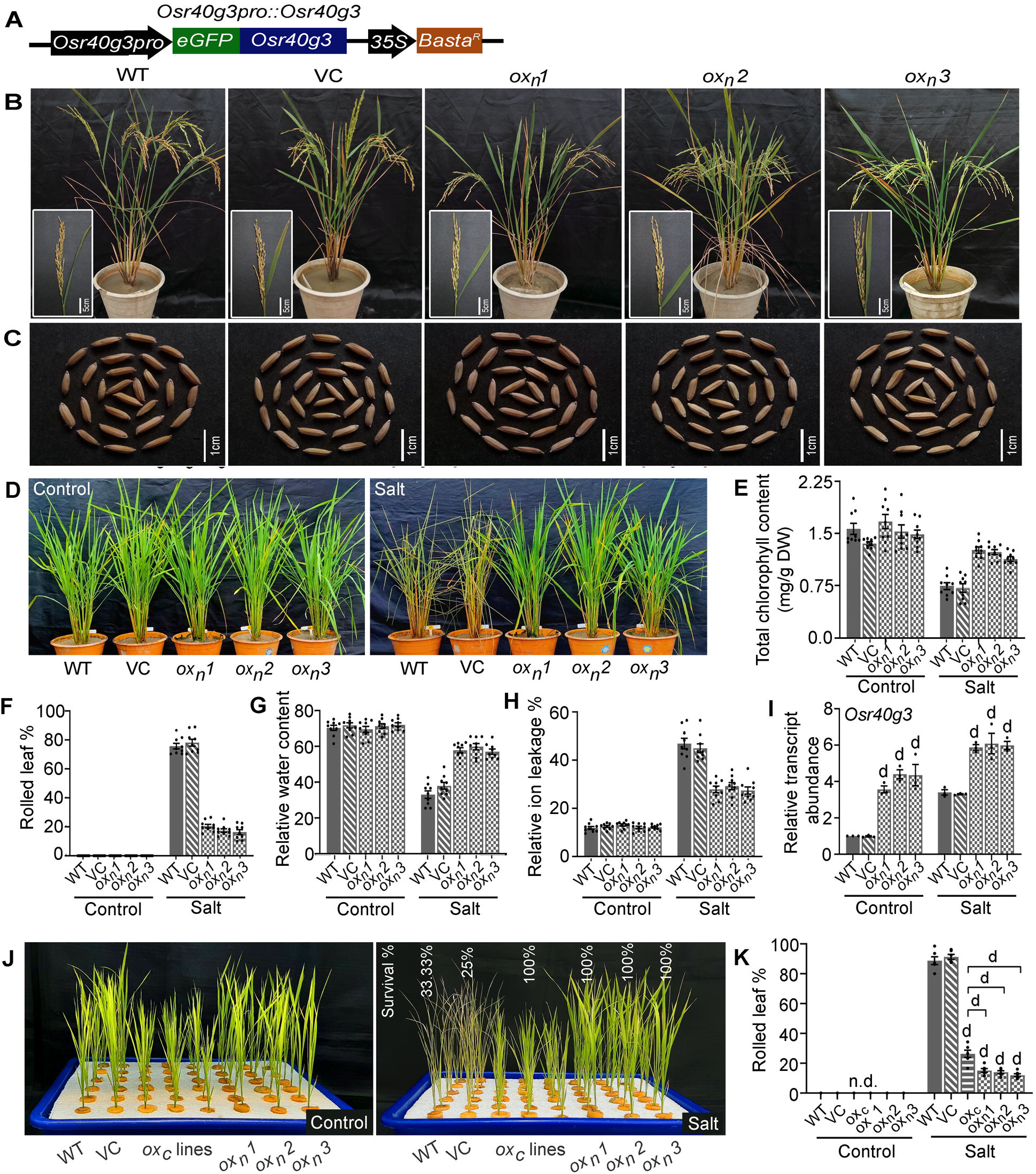
Overexpression of *Osr40g3* under control of its native promoter (*ox_n_* lines) improved salt tolerance with proper seed development. Different morphological parameters of 3 independent transgenic lines (*ox_n_*) harboring the *Osr40g3pro::Osr40g3* construct were recorded and compared with WT and VC (*35S::GFP*) plants. The WT, VC and 3 independent transgenic *ox_n_* lines were exposed to salt stress for 5 d and analyzed. (A) Overexpression construct, (B) WT, VC and transgenic *ox_n_* lines at reproductive stage with panicles in the inset, (C) grain morphology, (D) morphological response of WT, VC and 3 independent *ox_n_* lines under salt stress, (E) total chlorophyll content, (F) rolled leaf percentage, (G) relative water content, (H) relative ion leakage percentage, and (I) relative transcript abundance of *Osr40g3* gene in response to salt stress. For morphological analysis, 10 biological replicates were considered for each line (T_1_). The stress assay was independently repeated thrice with 3 biological replicates (T_1_ lines) in each set. Salt tolerance potential of the *ox_c_*, and *ox_n_* lines were compared and survival percentage was analyzed. (J) Morphological response of hydroponically grown WT, VC and independent *ox_c_*, and *ox_n_* lines under salt stress, and (I) rolled leaf percentage. For stress assay, 6 biological replicates for each of WT, VC, and 3 independent *ox_n_* lines (T_1_) were considered whereas 18 independent *ox_c_* lines (T_0_) were used. Results were represented as mean ± SEM and the statistical difference between WT, VC and transgenic lines was denoted by ‘d’ at P<0.0001. n.d.: not detected.

### *Os*r40g3 positively regulated *Os*EG45 protein while hampering the stability of *Os*GF14e

To gain mechanistic insight into the function of *Osr40g3*, we identified its interacting protein partners by yeast two-hybrid (Y2H) analysis against specific cDNA libraries. When we used a cDNA library from salt-stressed rice plants, the *Os*EG45-like domain-containing protein (*Os*EG45; LOC4347346), a peroxisomal membrane protein, *Os*PMP22 (LOC4346360), and 60S ribosomal protein L35 (LOC4329423) were identified as interacting partners (Figure 5A). Out of these partners, *OsEG45*, an expansin-like protein, was up-regulated in response to salt stress (Figure 5B). Again, *Os*r40g3 interacted with three non-redundant proteins, namely *Os*GF14e (LOC4329775), *Os*AGL20 (LOCOs03g03070), and *Os*AP2-like protein (LOCOs0107120), when a cDNA library prepared from rice flowers was used (Figure 5C). Among these partners, *Os*GF14e protein is associated with seed development in rice (Manosalva *et al*., 2011; Liu *et al*., 2015). We, therefore, selected *Os*EG45 and *Os*GF14e for further analysis and confirmed their interaction with *Os*r40g3 by bimolecular fluorescence complementation (BiFC) and co-immunoprecipitation (Co-IP) assays (Figure 5D-F).

**Figure 5.**
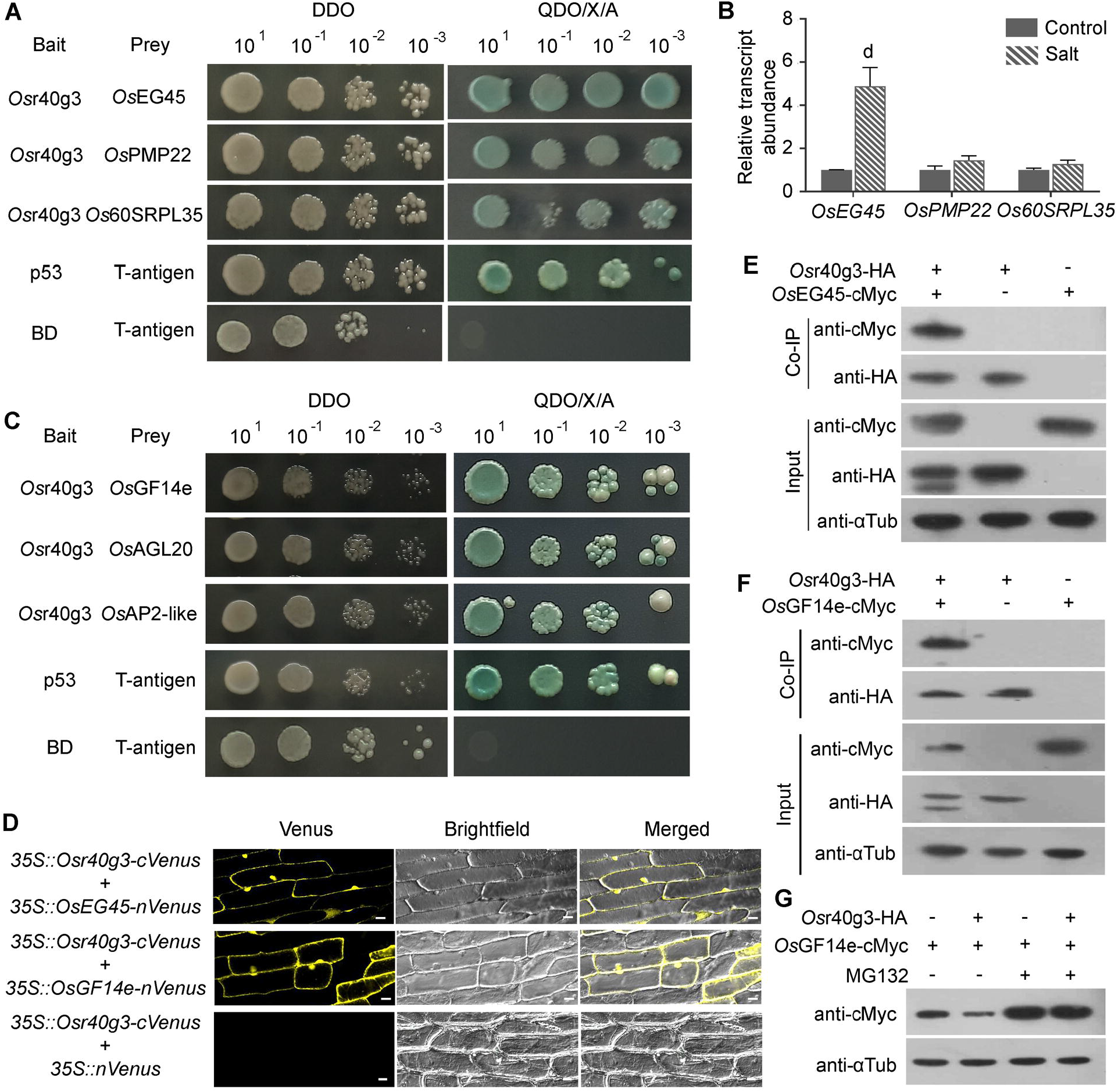
Identification of interacting protein partners of *Os*r40g3. (A) Yeast two-hybrid assay identified the interaction of *Os*r40g3 with *Os*EG45-like domain containing protein (*Os*EG45), a peroxisomal membrane protein, *Os*PMP22 and 60S ribosomal protein L35 when a cDNA library from rice leaves under salt stress was used. The interaction of p53 protein with T-antigen was used as a positive control. The interaction of the empty BD with T-antigen was used as a negative control. (B) Relative transcript abundance of the identified partners in response to salt stress. *OsEG45* was found to be salt-responsive. Results were represented as mean ± SEM and the statistical difference between control and salt conditions was denoted by ‘d’ at P<0.0001. (C) When a cDNA library from rice flowers was used, yeast two-hybrid analysis identified the interaction of *Os*r40g3 with *Os*GF14e, *Os*AGL20 and *Os*AP2-like proteins. The interaction of p53 protein with T-antigen was used as a positive control. The interaction of the empty BD with T-antigen was used as a negative control. (D) BiFC analysis confirmed the interaction of *Os*r40g3 protein with *Os*EG45 and *Os*GF14e. Venus fluorescence, bright field, and merged images were represented for each set of constructs. (E) Co-immunoprecipitation analysis was performed for the interaction of *Os*r40g3-HA with *Os*EG45-cMyc followed by immunoblot analysis using anti-HA and anti-cMyc antibody. The experiment was independently repeated thrice. (F) Interaction of *Os*GF14e-cMyc with *Os*r40g3-HA was similarly confirmed through co-immunoiprecipitation assay followed by immunoblot analysis using anti-HA and anti-cMyc antibody. The experiment was independently repeated thrice. (G) Proteasomal degradation assay of *Os*GF14e-cMyc was performed using a proteasomal inhibitor, MG132 followed by immunoblot analysis using anti-cMyc antibody. The amount of *Os*GF14e-cMyc was reduced to a significant level when co-expressed with *Os*r40g3-HA. The experiment was independently repeated thrice. Immunoblot analysis using anti-*α* tubulin antibody against total protein was used as loading control.

Intriguingly, when we performed immunoblot analysis from total protein against anti-cMyc antibody, significantly lower band intensity was observed in the case of calluses co-expressing both *Os*r40g3-HA and *Os*GF14e-cMyc proteins (Figure 5F). However, we did not detect any differential band intensity for the *Os*r40g3-HA and *Os*EG45-cMyc combination (Figure 5E). This data prompted us to explore if the binding of *Os*r40g3 triggers *Os*GF14e protein degradation. To investigate this possibility, we treated the transfected rice calluses with a proteasomal degradation inhibitor, MG132. When we performed immunoblot analysis using an anti-cMyc antibody, the band intensity was lower in the case of calluses co-transfected with *Os*r40g3-HA and *Os*GF14e-cMyc. In contrast, when we treated the co-transfected calluses with MG132, the band intensity remained unaltered. This observation indicated the proteasomal degradation of *Os*GF14e protein in presence of *Os*r40g3 (Figure 5G). In line with this observation, *OsGF14e* expression was higher in flowers and stamens of the WT plants with lower transcript abundance in leaves and roots (Figure S13). *Osr40g3*, on the contrary, followed an opposite expression pattern.

Although *Arabidopsis* has several GF14 orthologs, *Osr40g3* ectopic expression did not interfere with its seed development. Therefore, we analyzed the interaction of all the GF14 proteins in *Arabidopsis* with *Os*r40g3 through Y2H analysis, but no interaction was evident (Figure S14). On the other hand, the expansin-like B2 precursor protein (*At*EXLB2), an ortholog of *Os*EG45, was found to interact with the *Os*r40g3 protein. Multiple sequence alignment analysis revealed a highly conserved nature of the GF14 proteins except at their C-terminal domains (Figure S15). This structural distinctiveness of the *Os*GF14e presumably accounted for its binding specificity with *Os*r40g3, while its *Arabidopsis* orthologs failed to do so. Supporting this hypothesis, Y2H analysis confirmed that the C-terminal domain of *Os*GF14e is responsible for its interaction with *Os*r40g3 (Figure S16). In addition, the other members of the *Os*R40 family, like *Os*r40c1, *Os*r40g2, *Os*r40c2, and putative *Os*r40c1, did not interact with *Os*GF14e (Figure S17).

### *OsEG45* positively regulated salt tolerance

To validate the role of *Os*EG45 in regulating salt tolerance, we generated *OsEG45* overexpression (*OsEG45_ox_*) and silencing lines (*OsEG45_si_*). 14 independent transgenic lines were raised for *OsEG45_ox_* and *OsEG45_si_*, each. The transgenic plants displayed no phenotypic abnormalities during vegetative and reproductive stages. In response to salt stress, the *OsEG45_ox_* lines exhibited significantly improved salt tolerance over the WT and VC plants, while the *OsEG45_si_* lines showed increased sensitivity (Figure 6A-E, S18). The relative transcript abundance of the *OsEG45* gene was significantly higher in the *OsEG45_ox_* lines while lower in the *OsEG45_si_* lines compared with the WT and VC plants, thus confirming its crucial role in regulating salt tolerance (Figure 6F).

**Figure 6.**
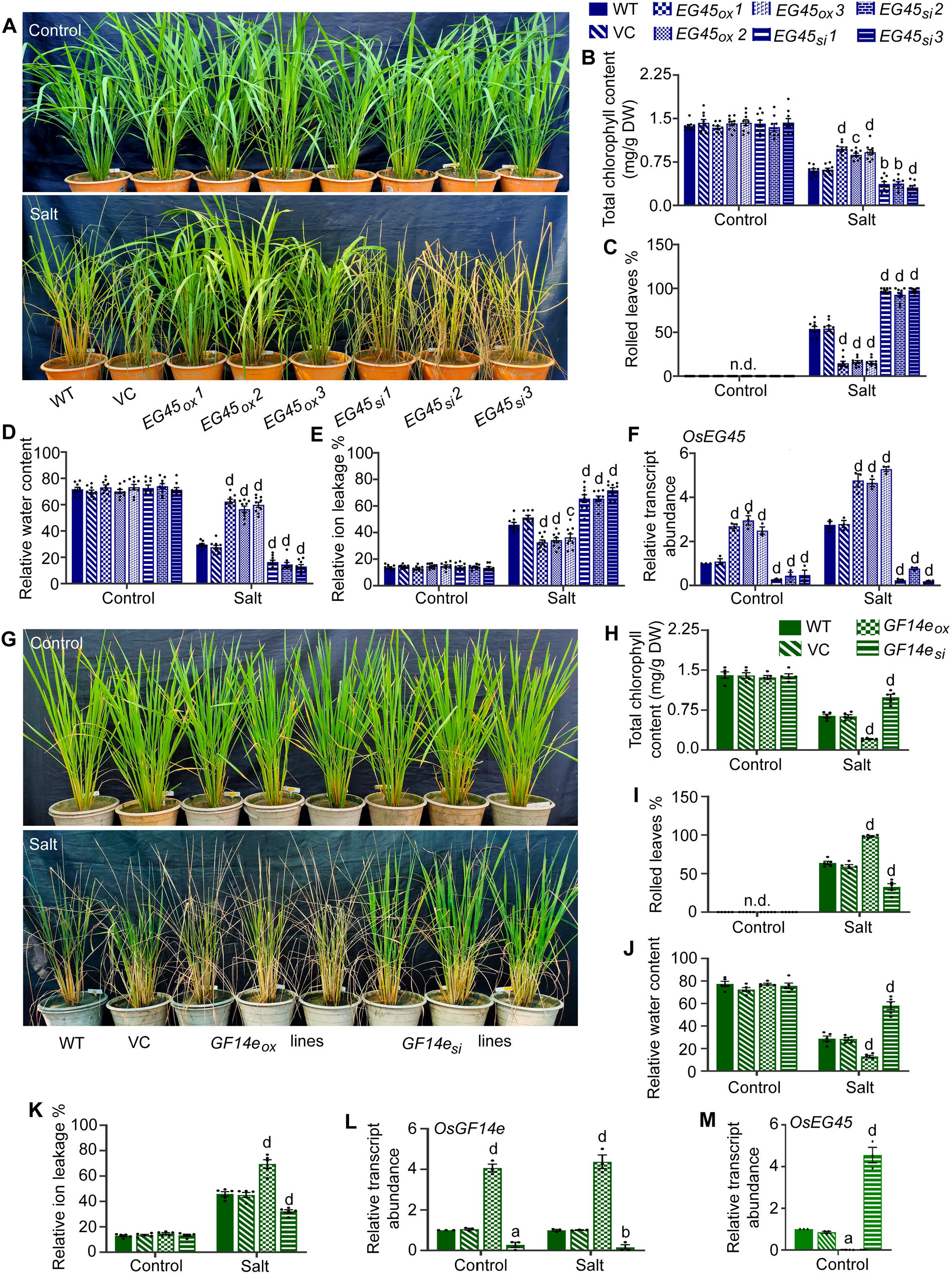
*Os*EG45 plays a positive role in regulating salt tolerance while *Os*GF14e plays a negative role. The WT, VC (*35S::GFP*) and 3 independent transgenic lines for each of the *OsEG45* overexpressing (*OsEG45_ox_*) and silencing (*OsEG45_si_*) lines were exposed to salt stress for 5 d and analyzed. (A) Morphological response of WT, VC and 3 independent *OsEG45_ox_* and *OsEG45_si_* lines under salt stress, (B) total chlorophyll content, (C) rolled leaf percentage, (D) relative water content, (E) relative ion leakage percentage, and (F) relative transcript abundance of *OsEG45* gene in response to salt stress. The stress assay was independently repeated thrice with 3 biological replicates in each set with T_1_ plants. The WT, VC and independent transgenic lines for each of the *OsGF14e* overexpressing (*OsGF14_ox_*) and silencing (*OsGF14_si_*) lines were exposed to salt stress for 5 d and analyzed. (G) Morphological response of WT, VC and representative 3 independent *OsGF14e_ox_* and *OsGF14e_si_* lines under salt stress, (H) total chlorophyll content, (I) rolled leaf percentage, (J) relative water content, (K) relative ion leakage percentage, and (L) relative transcript abundance of *OsGF14e* gene in response to salt stress. The stress assay was independently repeated twice with 5 independent *OsGF14_ox_* and *OsGF14_si_* lines (T_0_). (M) The relative transcript abundance of *OsEG45* gene was analyzed in 3 independent *OsGF14_ox_* and *OsGF14_si_* lines. Results were represented as mean ± SEM and the statistical difference between WT, VC and transgenic lines was denoted by different letters at P<0.05 (a), P<0.01 (b), P<0.001 (c), and P<0.0001 (d). n.d.: not detected.

### Silencing of *OsGF14e* induced pollen sterility but improved salt tolerance

Next, to understand the role of *Os*GF14e in regulating seed development, we developed 13 and 12 independent *OsGF14e* overexpressing (*OsGF14e_ox_*) and silencing *OsGF14e_si_*) lines respectively and recorded different morphological parameters. Although we observed no abnormalities at the vegetative stage, the *OsGF14e_si_* lines failed to produce seeds, while seed development in the *OsGF14e_ox_* lines was not compromised (Figure S19). Among the 12 *OsGF14e_si_* lines generated, four lines developed only 18-27 seeds per plant. Pollen grains of *OsGF14e_si_* lines displayed severe sterility, while *OsGF14e_ox_* lines showed fertile pollen grains (Figure S20). This observation demonstrated a positive role of *Os*GF14e in regulating pollen fertility in rice.

To further dissect any possible salt-responsive role of *Os*GF14e, the *OsGF14e_ox_* and *OsGF14e_si_* lines were exposed to salt stress and analyzed. Intriguingly, the *OsGF14e_si_* lines exhibited improved salt tolerance, while the *OsGF14e_ox_* lines displayed severe sensitivity (Figure 6G-K, S21). The negative role of *Os*GF14e in regulating salt tolerance made us curious if it also regulates *Os*EG45. The relative transcript abundance of *OsEG45* was indeed found to be higher in *OsGF14e_si_* lines and lower in *OsGF14e_ox_* lines as compared with the WT and VC lines suggesting that the salt tolerance of *OsGF14e_si_* plants is possibly due to an increased expression of *OsEG45* (Figure 6L-M). In line with this observation, the *OsGF14e_si_OsEG45_si_* double mutant lines displayed severe salt sensitivity, indicating that the salt tolerance in the *OsGF14e_si_* lines is mediated through an induction of *OsEG45* (Figure S22). Since *Os*r40g3 does not interact with the *Os*GF14e orthologs in *Arabidopsis*, the expression of the *OsEG45* ortholog, *AtEXLB2*, in the *ec* lines was similar to the WT and VC plants under control and salt stress conditions (Figure S23). Next, we wondered if *Os*GF14e can regulate transcriptional activation of *Osr40g3*. Therefore, its possible interaction with the *OsEG45* promoter was analyzed using a yeast one-hybrid (Y1H) assay. However, no interaction was detected between them, suggesting an involvement of some other regulatory mechanism (Figure S24).

### Lower *Os*GF14e abundance reduced GA accumulation and imparted salt tolerance in rice

The germination efficiency of the few seeds developed in the *ox_c_* and *OsGF14e_si_* lines were analyzed and compared with the WT, VC, *ox_n_*, and *OsGF14e_ox_* lines. We observed that none of the seeds from the *ox_c_* and *OsGF14e_si_* lines germinated, while the germination percentage for the other lines was more than 90 % (Figure 7A-D). Since GA regulates dwarfism, pollen sterility, and seed germination, we were curious if the GA pathway could mediate the phenotypic abnormalities in these lines. Indeed, exogenous GA_3_ treatment resulted in 90-95 % seed germination in the otherwise dormant *ox_c_* and *OsGF14e_si_* seeds. In addition, the α-amylase activity was significantly lower in the *ox_c_* and *OsGF14e_si_* lines, which increased in response to exogenous GA_3_ treatment (Figure 7B, 7D). Further, we measured the GA_3_ content from *ox_c_, ox_n_*, and *OsGF14e_si_* lines. Intriguingly, GA_3_ content was significantly lower (undetectable) in both *ox_c_* and *OsGF14e_si_* lines compared to the WT, with no significant alteration in *ox_n_* lines (Figure 7E).

**Figure 7.**
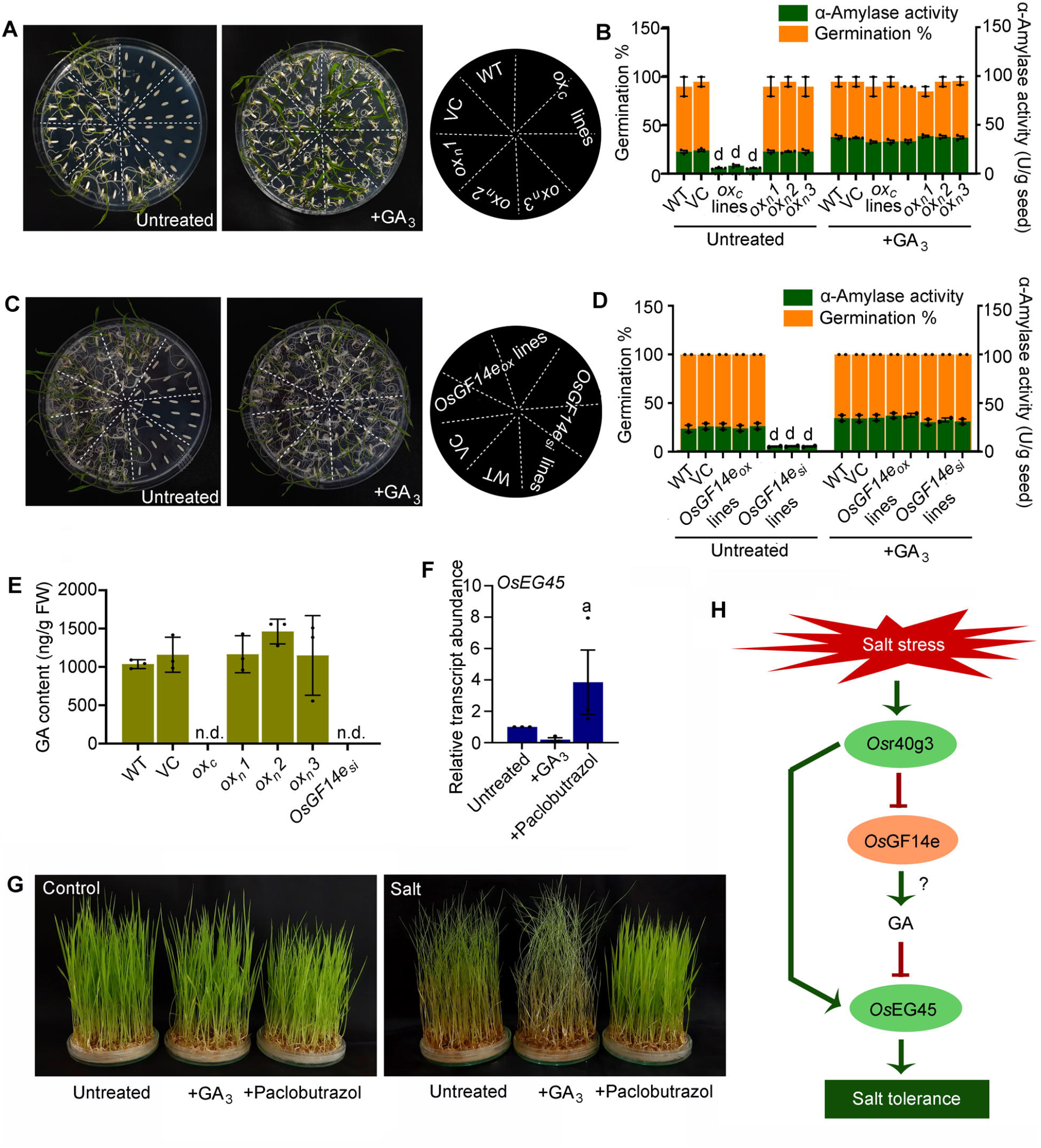
Constitutive overexpression of *Osr40g3* and silencing of *OsGF14e* interfered with GA metabolism in rice. (A) Seed germination assay for WT, VC (*35S::GUS*) and transgenic rice lines harboring *35S::Osr40g3 (ox_c_*) and *Osr40g3pro::Osr40g3 (ox_n_*) constructs under control and GA_3_ treated conditions. The experiment was independently repeated twice with 10 seeds of different independent lines. (B) Seed germination percentage and α-amylase assay for the same. The α-amylase assay was repeated thrice. (C) Seed germination assay for WT, VC, *OsGF14e_ox_* and *OsGF14e_si_* under control and GA_3_ treated conditions. The experiment was performed once since very few seeds were produced in the *OsGF14e_si_* lines. (D) Seed germination percentage and α-amylase assay for the same. The α-amylase assay was repeated twice. (E) The GA_3_ content was analyzed from WT, VC, *ox_c_, ox_n_*, and *OsGF14e_si_* lines. The experiment was independently repeated thrice. (F) 15-days-old WT seedlings were treated with exogenous GA_3_ or paclobutrazol for 48 h and the relative transcript abundance of *OsEG45* was analyzed. (G) Salt tolerance potential of the GA_3_ or paclobutrazol treated WT seedlings were analyzed. The experiment was independently repeated thrice. The results were represented as mean ± SEM. Statistical difference of the samples was denoted by different letters at P<0.05 (a), and P<0.0001 (d). n.d.: not detected. (H) Proposed model for *Os*r40g3-*Os*GF14e-*Os*EG45 module in regulating salt tolerance in rice.

Moreover, we were curious about any relationship between the reduced GA content and a higher abundance of *OsEG45* transcripts in the *OsGF14e_si_* plants. To investigate this, we analyzed the expression of *OsEG45* under altered GA conditions. Excitingly, when we treated rice seedlings with exogenous GA_3_, the transcript abundance of *OsEG45* was significantly reduced while it increased in response to the GA biosynthesis inhibitor, paclobutrazol (Figure 7F). In line with this observation, the rice seedlings displayed increased salt sensitivity in response to exogenous GA_3_ treatment while paclobutrazol treatment improved their salt tolerance (Figure 7G).

### A precise spatiotemporal regulation of the *Osr40g3-OsGF1e-OsEG45* module orchestrated salt tolerance in rice

Since *Osr40g3* and *OsGF14e* displayed a precise tissue-specific expression pattern, we also analyzed the relative transcript abundance of *OsEG45* in the root, leaf, and flower of the WT plant. Similar to *Osr40g3*, the expression of *OsEG45* was highest in the leaf (Figure S25). Next, we were curious if the regulation also involves temporal separation of *Osr40g3, OsEG45*, and *OsGF14e* gene expression in the leaves under salt stress. We, therefore, analyzed the relative transcript abundance of these three genes on a temporal landscape in response to salt stress. Interestingly, the *OsGF14e* expression was induced within 2 h of salt treatment, with the highest expression at 4 h. The expression then gradually declined within 12 h of salt stress. On the contrary, both *Osr40g3* and *OsEG45* displayed a late response to salt stress with the highest expression level at 5 d of salt treatment (Figure S26). This observation suggested that precise spatiotemporal regulation of these three genes modulates salt tolerance in rice.

## DISCUSSION

The osmotic stress-responsive nature of the R40 family proteins is well-described in plants (Moons *et al*., 1995; 1997). Among them, *Os*r40c1 plays a fundamental role in imparting drought tolerance in rice (Sahid *et al*., 2020). However, the functional aspects of the other *Os*R40 proteins remained unexplored to date. In the present investigation, we observed that among all the *Os*R40 family members, *Osr40g3* displayed the highest expression in response to salt stress (Figure S3). It prompted us to explore how *Osr40g3* regulates salt response in plants. Surprisingly, constitutive overexpression of the *Osr40g3* gene enhanced salt tolerance but resulted in pollen sterility in rice (Figure 1, 2, S4). On the contrary, the transgenic *Arabidopsis* lines ectopically expressing *Osr40g3* displayed improved salt tolerance without any defect in pollen viability and seed development (Figure S5-S7). The phenotypic abnormalities in the *ox_c_* lines motivated us to explore if the *Osr40g3* gene exhibits a tissue-specific expression pattern. Indeed, the gene displayed the highest expression in leaves with no expression in stamens (Figure 3). In line with this observation, overexpression of *Osr40g3* under the control of the native promoter significantly improved salt tolerance and displayed normal seed development, thus rescuing the pollen-sterile phenotype (Figure 4, S9, S10, S11). These observations indicated that *Osr40g3* follows a precise tissue-specific expression pattern, which is crucial for proper pollen viability and seed development in rice. Furthermore, several independent T_1_ lines were used for the analysis which might include a mix of homozygous and heterozygous lines. However, the lines displayed homogeneity in their stress response indicating that the heterozygous lines are as tolerant as its homozygous counterpart.

Identifying the interacting protein partners of *Os*r40g3 was the next relevant point in solving the functional mechanism of the protein. *Os*r40g3 interacted with the *Os*EG45 protein in the salt-stressed rice leaves (Figure 5). EG45 is a salt stress-responsive expansin-like glycoprotein that shows endoglucanase-like function and helps in cell wall modification (Ludidi *et al*., 2002). Cell wall modification is a common phenomenon observed in osmotically stressed cells (Gall *et al*., 2015). Several expansin genes are known to be up-regulated under osmotic stress. For example, the rose expansin gene, *RhEXPA4*, is induced in response to salt stress, and its ectopic expression in *Arabidopsis* leads to a salt-tolerant phenotype (Lü *et al*., 2013). Likewise, transgenic tobacco expressing wheat expansin gene *TaEXPB23* displays improved salt tolerance (Han *et al*., 2012). In the present investigation, overexpression of *OsEG45* led to improved salt tolerance, while silencing of this gene displayed sensitivity towards salt stress (Figure 6). This observation suggested that *Os*r40g3 significantly improved salt tolerance in rice by positively regulating *Os*EG45 protein. Since *Os*r40g3 protein could also interact with *At*EXLB2, the *Arabidopsis* ortholog of *Os*EG45, the *ec* lines displayed salt tolerance (Figure S14).

On the other hand, when we used a cDNA library from rice flowers, *Os*r40g3 interacted with *Os*GF14e (Figure 5). *Os*GF14e is a member of the GF14 protein family (14-3-3 proteins) and modulates several stress responses and developmental phenomena in plants (Chen *et al*., 2006). For example, the rice GF14 family proteins like *Os*GF14b, *Os*GF14c, *Os*GF14e, and *Os*GF14f interact with different target proteins to critically regulate signaling responses during biotic stress (Chen *et al*., 2006). Furthermore, *Os*GF14e negatively regulates sheath blight and bacterial blight resistance in rice (Manosalva *et al*., 2011). Also, suppression of *Os*GF14e could produce susceptibility toward panicle blast (Manosalva *et al*., 2011; Liu *et al*., 2015). Earlier evidence suggested that low accumulation of GF14e protein is associated with sterile pollen grains in maize (Datta *et al*., 2002). Again, the silencing of the *OsGF14e* gene displays stunted growth and poor seed development in rice (Manosalva *et al*., 2011). A subsequent study by Liu *et al*. (2015) precisely demonstrated that all the *OsGF14e*-silenced lines failed to develop seeds except for two lines with less *OsGF14e* silencing. Similarly, in the present study, *Os*r40g3 was found to negatively regulate the stability of *Os*GF14e protein, which presumably led to impaired seed development in the *ox_c_* lines (Figure 1, 5). Further, the *OsGF14e_si_* lines also displayed pollen sterility and failed to produce seeds, which corroborated with the previous reports (Figure S19, S20). Accumulating evidence also suggested that rice GF14 family members interact with site-specific DNA binding proteins like *EmBP1* and tissue-specific transcription factor VP1 to regulate signaling responses in a spatial manner (Schultz *et al*., 1998). It supports the precise tissue-specific function of the GF14 proteins in plants. Consistent with these reports, out of the eight GF14 family members, the *Os*GF14e and *Os*GF14f show higher accumulation in flowers and panicles, which corroborates our present observation (Chen *et al*., 2006).

Furthermore, a few seeds that developed in the *ox_c_* and *OsGF14e_si_* lines displayed complete inhibition of germination. Since dwarfism, pollen sterility, and seed germination are widely known to be regulated by GA, we assumed that these transgenic lines might have a defect in GA metabolism. Indeed, exogenous application of GA_3_ could rescue the germination inhibition in the transgenic seeds supporting this hypothesis (Figure 7). The lower accumulation of GA_3_ in these transgenic lines further confirmed the alteration in GA metabolism (Figure 7). These observations corroborated with the previous report where *OsGF14e*-silenced rice plants also exhibited a defect in GA metabolism (Liu *et al*., 2015). However, how *Os*GF14e integrates the GA pathway with pollen development is an exciting area to explore in the future.

Although *Os*GF14e has several orthologs in *Arabidopsis*, the *ec* lines displayed normal seed development. To understand this unusual observation, we checked if these orthologs in *Arabidopsis* could interact with *Os*r40g3. However, none of them could interact with *Os*r40g3 suggesting high specificity of the *Os*r40g3-*Os*GF14e interaction (Figure S14). Despite their high sequence homology, the varied C-terminal domain of the GF14 family members determines their functional variation and specificity in plants (Sehnke *et al*., 2006). Indeed, only the C-terminal domain of *Os*GF14e is responsible for its interaction with *Os*r40g3 (Figure S16). This structural distinctiveness of *Os*GF14e from its *Arabidopsis* orthologs provides a possible answer to why *Os*r40g3 ectopic expression failed to impede seed development in *Arabidopsis*.

The *Os*r40g3 interacted with *Os*GF14e in stamens only when overexpressed under a constitutive promoter. In the WT, however, the *Os*r40g3 is not expressed in stamen, ruling out its scope of interaction with *Os*GF14e. We, therefore, wondered why this specific interaction evolved and if it has any role in modulating *Os*r40g3-mediated salt stress tolerance in rice. The higher transcript abundance of *OsEG45* in the salt-tolerant *OsGF14e_si_* lines and its lower abundance in the salt-sensitive *OsGF14e_ox_* lines suggested intricate signaling crosstalk between *Os*r40g3, *Os*GF14e, and *Os*EG45 in regulating salt tolerance in rice (Figure 6). Moreover, in the leaf, the expression maxima of *OsGF14e* and *Osr40g3* are separated temporally during salt stress (Figure S26). This observation corroborates the previous report where *OsGF14e* expression rapidly increased in response to salt exposure and then sharply declined after 8 h in rice (Chen et al., 2006). Another study demonstrated that, following dehydration stress, endogenous GA levels increase between 3-6 h and then decline after 24 h (Urano *et al*., 2017). On the contrary, the *OsEG45* expression peaked after 5 d of salt stress which overlapped with the *Osr40g3* expression maxima (Figure S26). We, therefore, hypothesized that during salt stress, the *Os*r40g3-mediated proteasomal degradation of *Os*GF14e results in a lower accumulation of endogenous GA in plants which activates *OsEG45* expression leading to salt tolerance. Supporting this hypothesis, silencing the *OsEG45* gene in *OsGF14e_si_* lines enhanced salt sensitivity, thus elucidating an *Os*GF14e-GA-*Os*EG45 mediated regulation of salt stress response in rice. Again, the exogenous GA_3_ treatment enhances the salt sensitivity in rice and diminishes *OsEG45* gene expression, further supporting our hypothesis (Figure 7). In line with this observation, earlier reports also documented that lowering GA accumulation improves salt tolerance in rice (Shan *et al*., 2014; Zhou *et al*., 2020; Yu *et al*., 2020). The diminished GA accumulation leading to growth retardation under salt stress represents an adaptive strategy for survival.

In summary, we conclude that *Os*r40g3 negatively regulates *Os*GF14e resulting in perturbation of GA metabolism. The reduced GA content, in turn, induces the expression of *OsEG45*. In addition, *Os*r40g3 positively regulates *Os*EG45 by directly interacting with it (Figure 7H). Together the present investigation unraveled a novel *Os*r40g3-*Os*GF14e-*Os*EG45 signaling module in mitigating salt stress that might be exploited for developing salt-tolerant rice in future crop improvement programs.

## EXPERIMENTAL PROCEDURES

### Plant material and stress treatment

The seeds of different *indica* rice cultivars like Nonabokhra, Ranjit, Khitish, IR64, Swarna sub1, Jaldi13, MTU1010, and Vandana were procured from Chinsurah Rice Research Station, West Bengal, India. Plants were grown in a greenhouse under the optimum condition at a day/night temperature regime of 30°-35°C/22°-29°C and 60-65 % relative humidity with 12 h photoperiod. For salt stress, seeds were surface sterilized using 4 % sodium hypochlorite followed by 0.2 % bavistin treatment, germinated, and grown on filter papers wetted with water for 15 d. A hundred seedlings of each cultivar were treated with 200 mM NaCl solution for 5 d as standardized before (Kurniasih *et al*., 2013; Sahid *et al*., 2020). For the control set, an equal volume of distilled water was used. Following stress treatment the cultivars were assessed according to the standard evaluation system proposed by IRRI (2013). For salt stress treatment of soil-grown plants, the 60-days-old plants were irrigated with 200 mL of 200 mM NaCl solution per day, for 5 consecutive days, and data was recorded after 5 d of salt treatment (Amirjani, 2012). Five leaves were harvested from each plant on the 0^th^ day of stress treatment as control sample and analyzed. The Na^+^, K^+^, and Cl^-^ content as well as EC of the soil were estimated before and after salt treatment (Khan and Amin, 2019). For salt stress treatment of hydroponically grown plants, 30-days-old plants grown in Yoshida medium were used (Yoshida *et al*., 1976). The plants were then exposed to Yoshida medium containing NaCl at a final concentration of 200 mM for 5 d and analyzed.

*Arabidopsis thaliana* seeds of Columbia ecotype (Col-0) were procured from Nottingham Arabidopsis Stock Center (NASC), UK, and were grown in a growth chamber at 22°C under 16 h light/8 h dark cycle as standardized before (Datta *et al*., 2015). 30-days-old plants were exposed to salt stress by irrigating with 20 mL of 200 mM NaCl solution every alternate day as standardized before (Datta *et al*., 2015). Five leaves were harvested from each plant on the 0^th^ day of stress treatment as control sample and analyzed.

### Morphological analyses

Different morphological parameters like shoot and root length, seedling dry weight, and percentage of rolled leaves were analyzed under control and stressed conditions. The tissue was dried at 50°C for 3 d to measure the dry weight. The RWC was estimated after salt stress exposure following Ma *et al*. (2018).

### Estimation of different biochemical compounds

Proline and glycine betaine contents were measured by spectrophotometric method from tissue samples collected from control and treated plants according to Grieve and Grattan (1983) and Woodrow *et al*. (2016) respectively. The chlorophyll content was measured according to Lichtenthaler (1987). The H_2_O_2_ content was measured following Yin *et al*. (2010).

### RNA extraction and qRT PCR analysis

Total RNA was extracted using the Trizol method and cDNA was synthesized using the iScript™ cDNA Synthesis Kit (Bio-Rad). qRT PCR analysis was performed in CFX96 Touch™ Real-Time PCR Detection System (Bio-Rad) using iTaq Universal SYBR Green Supermix (Bio-Rad) and gene-specific primers (Supplemental Table S2). Gene expression was normalized based on the relative quantification method. The *Actin* gene of rice and *Arabidopsis* was used as a reference gene.

### Vector construction and plant transformation

For the development of overexpression lines, the full-length gene of *Osr40g3* (LOC112936024) was amplified and cloned between the *EcoRI* and *BamHI* restriction enzyme (RE) sites of the *pEGAD* vector under the control of a constitutive *CaMV35S* promoter. The recombinant construct (*35S::Osr40g3*) was introduced into the background of *indica* rice cultivar khitish (IET4095) following *Agrobacterium*-mediated genetic transformation (Datta *et al*., 2000; Sahid *et al*., 2020). The putative transformants were screened for herbicide resistance, and positive lines were confirmed by genomic DNA PCR using *bar* gene-specific primers. The same construct was used to transform *Arabidopsis* plants of Col-0 ecotype via the *Agrobacterium*-mediated floral-dip method (Clough and Bent, 1998; Datta *et al*., 2015).

To clone the promoter region of the *Osr40g3* gene, the 1,315 bp upstream sequence from the +1 site of the *Osr40g3* gene was inserted between the *AgeI* and *PacI* RE sites of the recombinant vector *35S*::Os*r40g3* to generate the construct, *Osr40g3pro::Osr40g3*. The promoter fragment was also sub-cloned between the *BamHI* and *BglII* RE sites of the *pCAMBIA1304* vector to generate the *Osr40g3pro::GUS* construct. The constructs were used for *Agrobacterium*-mediated genetic transformation of rice as described above.

For the generation of the overexpression constructs for *OsEG45* and *OsGF14e* genes, the cds regions were cloned into *EcoRI* and *BamHI* RE sites of the *pEGAD* vector and the *SpeI* and *BamHI* sites of the *pVYNE* vector respectively. For the generation of silencing constructs for *OsEG45* and *OsGF14e* genes, the 266 bp and 335 bp coding sequences, respectively, were amplified and cloned into the *pRNAi-GG* vector by the Golden Gate cloning method following Yan *et al*. (2012). The constructs were used for *Agrobacterium*-mediated genetic transformation of rice as described above. For the generation of the double mutant, silencing constructs of *OsEG45* and *OsGF14e* were co-transformed into rice following the *Agrobacterium*-mediated genetic transformation method.

### Morphological analysis of transgenic plants

Morphological parameters like plant height, number of tillers, shoot and root dry weights, panicle length, number of panicles/plant, number of flowers/panicle, number of grains/panicle, and floral morphology were analyzed from the WT, VC, and transgenic rice plants. In the case of *Arabidopsis*, plant height, rosette diameter, floral morphology, silique length, number of silique/plant, number of seeds/silique, and seed yield/plant were measured from WT, VC, and transgenic lines. Floral morphology of both rice and *Arabidopsis* plants was studied and documented under a zoom stereo trinocular microscope (Magnus MSZ-RR LED). For salt stress assay, different morphological parameters were analyzed under control and stressed conditions as described above.

### Estimation of Na^+^ and K^+^ contents

The Na^+^ and K^+^ contents were measured by a flame photometer (Systronics, model 128) following Nublat *et al*. (2001) with slight modifications. Briefly, leaves were collected from plants under control and stress conditions and dried at 80° C for 24 h. The dried samples were then incubated in 0.1 M nitric acid for 30 min. The extract was filtered and analyzed.

### Estimation of lipid peroxidation and ion leakage percentage

The lipid peroxidation of shoots was analyzed by measuring malondialdehyde (MDA) content following Yin *et al*. (2020). The relative electrolyte leakage was determined by analyzing relative conductivity in solution using a conductivity meter (Systronics) following Cao *et al*. (2007).

### Pollen viability assay

For I_2_-KI staining, mature rice pollen grains were stained with 2 % I_2_-KI (w/v) solution and observed under a bright field microscope (Leica DM IL LED S-80) according to Chhun *et al*., (2007). For FDA staining, mature pollen grains were incubated in pollen viability solution [PVS, 0.001 % (w/v) fluorescein diacetate, 290 mM sucrose, 1 mM KNO_3_, 0.16 mM boric acid, 1.27 mM Ca(NO_3_)_2_] in the dark for 15 min at 28°C and centrifuged. The pellet was resuspended in PVS without FDA and observed under a fluorescence microscope (Leica DM IL LED S-80) using 488 nm excitation and 510 nm emission wavelengths (Muhlemann *et al*., 2018). For *Arabidopsis* pollen grains, Alexander staining was performed according to Schofta *et al*. (2011) and examined under a bright field microscope (Leica DM IL LED S-80). At least 500 pollen grains were considered for each sample.

### *In vitro* pollen germination assay

*In vitro* pollen germination assay was performed according to Li *et al*. (2018). Briefly, the mature pollen grains collected from rice flowers were spread onto pollen germination medium [pH 6.0, containing of 15 % (w/v) sucrose, 0.01 % (w/v) boric acid, 1 mM Ca(NO_3_)_2_, and 0.3 % (w/v) low melting agarose] and were incubated at 25°C under humid condition. After 4 h of incubation, pollen germination was analyzed and documented under a bright field microscope (Leica DM IL LED S-80).

For *in vitro* pollen germination of *Arabidopsis*, a solid pollen germination medium was used. The mature pollen grains were collected and spread on the pollen germination medium containing 0.01 % H_3_BO_4_ (w/v), 5 mM CaCl_2_, 5 mM KCl, 1 mM MgSO_4_, 10 % sucrose (w/v), and 0.1 % agarose (Boavida and McCormick, 2007). The pollen germination was examined under a bright field microscope (Leica DM IL LED S-80) after 6 h incubation.

### Hybridization of rice plants

The flowers of WT plants were emasculated and considered female parents whereas the transgenic *ox_c_* plants were considered male parents. In another set, the transgenic *ox_c_* plants were emasculated and considered as the female parent and hybridized with WT as the male parent. Hybridization was carried out as standardized before (Paul *et al*., 2014). In both cases, the number of hybrid seeds was determined.

### Histochemical GUS assay

Samples from different tissues under various developmental stages were collected from the transgenic rice lines harboring the *Osr40g3pro::GUS* construct. Samples were stained with GUS staining solution (50 mM sodium phosphate, pH 7.0, 10 mM EDTA, 0.5 mM K_3_[Fe(CN)_6_], 0.5 mM K_4_[Fe(CN)_6_], 0.1 % Triton X-100 (v/v), and 1 mM X-glucuronide) following Jefferson *et al*. (1987). The stained samples were visualized under a zoom stereo trinocular microscope (Magnus MSZ-RR LED).

### Y2H analysis

Y2H analysis was performed using the Matchmaker^®^ Gold Yeast Two-Hybrid System (Takara) following the manufacturer’s protocol. The *Osr40g3* gene was inserted into the *EcoRI* and *BamHI* RE sites of the bait vector, *pGBKT7* (Takara). *Os*r40C1, *Os*r40g2, *Os*r40c2 and putative *Os*r40c1 were also cloned into the bait vector pGBKT7. The cDNA library was prepared from rice flowers and ligated to the *pGADT7-Rec* vector using the Make Your Own “Mate and Plate” Library System (Takara). A second cDNA library was prepared from rice leaves under salt stress. The recombinant plasmids were co-transformed into yeast Y2H Gold strain (Takara). The transformed yeast strains were grown on DDO (SD/-Leu/-Trp) and QDO/X/A (SD/-Ade/-His/-Leu/-Trp/X-α-Gal/Aureobasidin-A) media and analyzed by sequencing following the manufacturer’s instructions to identify the interacting protein partners. To analyze the interaction of *Os*r40g3 with the orthologs of *Os*EG45 and *Os*GF14e in *Arabidopsis*, the *At*EXLB2 and 12 *At*GF14 proteins were cloned into the *pGADT7-Rec* vector and used as prey for the *BD-Osr40g3* bait. To identify domain-specific interaction of the *Os*GF14e, its N-terminal and C-terminal domains were cloned into the *pGADT7-Rec* vector and used as prey for the *BD-Osr40g3* bait. Y2H analysis was performed as described above. The *BD-p53* and *AD-T-antigen* interaction was used as a positive control. The empty *BD* and *AD-T-antigen* interaction was used as a negative control.

### BiFC assay

For BiFC assay, the *Osr40g3* gene without stop codon was inserted into the vector *pVYCE* between *ApaI* and *BamHI* RE sites to obtain the *35S::Osr40g3-cVenus* construct. The *OsGF14e* and *OsEG45* genes were cloned into *SpeI* and *BamHI* RE sites and *SpeI* and *XbaI* RE sites of the *pVYNE* vector to generate the *35S::OsGF14e-nVenus* and *35S::OsEG45-nVenus* constructs respectively (Waadt *et al*., 2008). The recombinant constructs were introduced into *Agrobacterium tumefaciens* strain GV3101 and used to infiltrate onion epidermal cells (Yang *et al*., 2014). The fluorescence signal for Venus protein was detected at the excitation wavelength of 514 nm using a confocal laser scanning microscope (Olympus FluoView™ FV1000). The *35S::Osr40g3-cVenus* construct with an empty *35S::nVenus* vector was considered as a negative control.

### Co-IP and immunoblot analysis

The rice embryo calluses transformed with *35S::Osr40g3-HA* and/or *35S::OsGF14e-cMyc* were used for total protein isolation (Chen *et al*., 2019). The samples were extracted in lysis buffer (20 mM Tris□Cl pH 7.5, 150 mM NaCl, 0.5 % Triton X□100 (v/v) and protease inhibitor, Thermo Scientific) at 4°C and centrifuged at 13,000 rpm for 30 min at 4°C. Supernatants were mixed with 20 μL anti-HA-agarose beads and incubated overnight following the manufacturer’s instructions (Pierce™ HA-Tag IP/Co-IP Kit, Thermo Fisher Scientific). The HA-tagged protein was eluted and quantified by the Bradford method. An equal amount of protein was used for immunoblot analysis using an anti-HA antibody (1:1000 dilution, Sigma) and anti-cMyc antibody (1:1000 dilution, Agrisera). Immunoblot against anti-α-Tubulin antibody (1:1000 dilution, Agrisera) was used as a loading control. The co-IP assay for *Os*r40g3-HA and *Os*EG45-cMyc interaction was similarly performed. For proteasomal degradation inhibitor assay, the transformed rice calluses were incubated in dark for 10 d at 37°C. The calluses were then incubated with 10 μM MG132 for 4 h. Proteins were isolated from the calluses and used for immunoblot analysis as described above (Garcia-Cano *et al*., 2014).

### Y1H analysis

To identify the interaction of *OsGF14e* protein with the *OsEG45* promoter, the 1707 bp upstream sequence from the +1 site of the *OsEG45* gene was cloned into the *KpnI* and *SacI* RE sites of the *pAbAi* vector (Takara). The *OsGF14e* gene was cloned into the *pGADT7-Rec* vector (Takara). The recombinant constructs were introduced into Y1H gold strain and Y1H assay was performed using Matchmaker^®^ Gold Yeast One-hybrid screening system (Takara) following the manufacturer’s protocol. The transformed yeast strains were grown on SD/-Leu and SD/-Leu/Aureobasidin-A media.

### Seed germination and α-amylase assay

Seeds were surface sterilized with 4 % sodium hypochlorite solution followed by 0.2 % bavistin treatment and placed on MS medium without or with 5 μM GA_3_ for germination in the dark for 7 d. Estimation of α-amylase activity was carried out following Shen *et al*. (2019).

### Estimation of endogenous GA

The GA_3_ content was measured using high-performance liquid chromatography following Liu *et al*. (2021). Briefly, the leaves were homogenized and extracted with pre-chilled 80% methanol for 20 h at 4°C. After centrifugation, the supernatant was dried using a vacuum concentrator. The samples were dissolved in 100 μL ultrapure methanol supplemented with 0.1 M glacial acetic acid. After filtration, the sample was analyzed using Waters HPLC system multi-dimensional liquid chromatography (model 2695) with a C18 analytical column. The flow rate was 1 ml min^−1^ and the column temperature was set at 30°C. The chromatogram was analyzed by Empower 3.0 software.

### Exogenous GA and paclobutrazol treatment

The 15-days-old seedlings were treated with a 5 μM GA_3_ or 2 μM paclobutrazol solutions for 48 h (Yang *et al*., 2004; Schmidt *et al*., 2014). Total RNA was extracted from seedlings and the expression of the *OsEG45* gene was analyzed through qRT-PCR. For salt stress assay, the GA_3_ or paclobutrazol-treated seedlings were exposed to 200 mM NaCl for 5 d.

### Multiple sequence alignment analysis

The sequences of *Arabidopsis* GF14 family proteins and *Os*GF14e were used for multiple sequence alignment using the PRALINE multiple sequence alignment tool (http://ibivu.cs.vu.nl/programs/pralinewww/).

### Statistical analysis

Statistical analysis was performed using GraphPad Prism version 8.3.0 software (GraphPad Software, San Diego, California USA). The variation of morphological, and biochemical parameters as well as the relative transcript abundance among genotypes and treatments were analyzed following two-way ANOVA followed by Sidak’s multiple comparison tests. To compare the morphological parameters between WT, VC, and transgenic lines, one-way ANOVA followed by Dunnett’s multiple comparison tests was performed. Details of replicates, sample size, and the significance levels of P-values were indicated in respective figure legends. The data were represented as mean ± standard error of the mean (SEM).

## ACCESSION NUMBERS

The accession numbers of all genes used in this study has been listed in the Supplemental Table S2.

## ACKNOWLEDGEMENT

This work has been supported by Department of Science and Technology and Biotechnology, Government of West Bengal, India [BT(Budget)/RD-29/2016]. We thank Central Instrumentation Facility of Department of Botany, University of Calcutta and Department of Botany, Dr. A. P. J. Abdul Kalam Government College as well as confocal microscopic facility of DBT-IPLS, Department of Biochemistry, University of Calcutta. We thank Prof. Jörg Kudla (University of Munster, Germany) for providing *pVYNE* and *pVYCE* vectors and Prof. Prabod Trivedi (CSIR-National Botanical Research Institute, India) for kindly sharing *Agrobacterium tumefaciens* GV3101 strain.

## AUTHOR CONTRIBUTIONS

RD and SP conceived and designed the original research plan; CR raised the transgenic lines, performed stress assays, morphological, biochemical, microscopic and expression analyses, Co-IP and immune blot analysis; SS generated the constructs and performed BiFC, and Y1H analyses; SS and DS performed Y2H analysis; RD and SP analyzed the data and wrote the manuscript.

## SUPPORTING INFORMATION

**Figure S1**. Salt stress assay of 8 *indica* rice cultivars.

**Figure S2.** Morphological and biochemical analyses of the 8 *indica* rice cultivars in response to salt stress.

**Figure S3.** Relative transcript abundance of 5 different *Os*R40 family genes from rice shoots in response to salt stress condition.

**Figure S4.** Biochemical analyses of the transgenic rice lines constitutively overexpressing the *Osr40g3* gene (*ox_c_* lines) in response to salt stress.

**Figure S5**. Morphological characterization of transgenic *Arabidopsis* lines ectopically expressing *Osr40g3*.

**Figure S6.** Response of transgenic *Arabidopsis* lines ectopically expressing *Osr40g3* gene under salt stress condition.

**Figure S7.** Floral morphology of transgenic *Arabidopsis* lines ectopically expressing *Osr40g3*.

**Figure S8.** Histochemical GUS assay for *Osr40g3* promoter activity in response to salt stress.

**Figure S9**. The transgenic rice plants overexpressing *Osr40g3* under control its native promoter (*ox_n_* lines) displayed no phenotypic abnormality.

**Figure S10.** Pollen viability of transgenic rice lines overexpressing *Osr40g3* under control of native promoter.

**Figure S11**. Biochemical analyses of the transgenic rice lines overexpressing the *Osr40g3* gene (*ox_n_* lines) under control of the native promoter in response to salt stress.

**Figure S12.** Relative transcript abundance of *OsR40* genes in WT, VC, *ox_c_* and *ox_n_* lines plant.

**Figure S13.** Relative transcript abundance of *OsGF14e* in leaf, root, flower and stamen of WT plant.

**Figure S14.** Yeast two-hybrid assay for the interaction of *Os*r40g3 with the *Arabidopsis* orthologs of *Os*GF14e and *Os*EG45 proteins.

**Figure S15.** Multiple sequence alignment of *Os*GF14e protein with *Arabidopsis* GF14 family members showing structural variability at the C-terminal end.

**Figure S16.** Yeast two-hybrid assay for the interaction of *Os*r40g3 with N-terminal and C-terminal domains of *Os*GF14e.

**Figure S17.** Yeast two-hybrid assay for the interaction of *Os*GF14e with all the *Os*r40 family proteins.

**Figure S18.** Biochemical analyses of the *OsEG45* overexpressing (*OsEG45_ox_* lines) and silencing (*OsEG45_si_* lines) transgenic rice lines in response to salt stress.

**Figure S19.** Morphological characterization of the *OsGF14e* overexpressing (*OsGF14e_ox_* lines) and silencing (*OsGF14e_si_* lines) transgenic rice lines.

**Figure S20**. Pollen viability of the *OsGF14e* overexpressing (*OsGF14e_ox_* lines) and silencing (*OsGF14e_si_* lines) transgenic rice lines.

**Figure S21**. Biochemical analyses of the *OsGF14e* overexpressing (*OsGF14e_ox_* lines) and silencing (*OsGF14e_si_* lines) transgenic rice lines in response to salt stress.

**Figure S22.** Salt stress assay of the *OsGF14e_si_OsEG45_si_* double mutant.

**Figure S23.** Relative transcript abundance of *AtEXLB2* in WT, VC, and *ec* lines.

**Figure S24.** Yeast one-hybrid assay for the interaction of *Os*GF14e with the *OsEG45* promoter.

**Figure S25.** Relative transcript abundance of *OsEG45* in different tissues of WT rice.

**Figure S26.** Expression analysis of *Osr40g3, OsGF14e*, and *OsEG45* genes from leaves in response to salt stress on a temporal landscape.

**Table S1.** Estimation of soil salt concentration and EC before and after salt stress

**Table S2.** Primers used in this study

