## Supplementary Figure for "*Os*r40g3 imparts salt tolerance by regulating GF14e-mediated gibberellin metabolism to activate EG45 in rice"

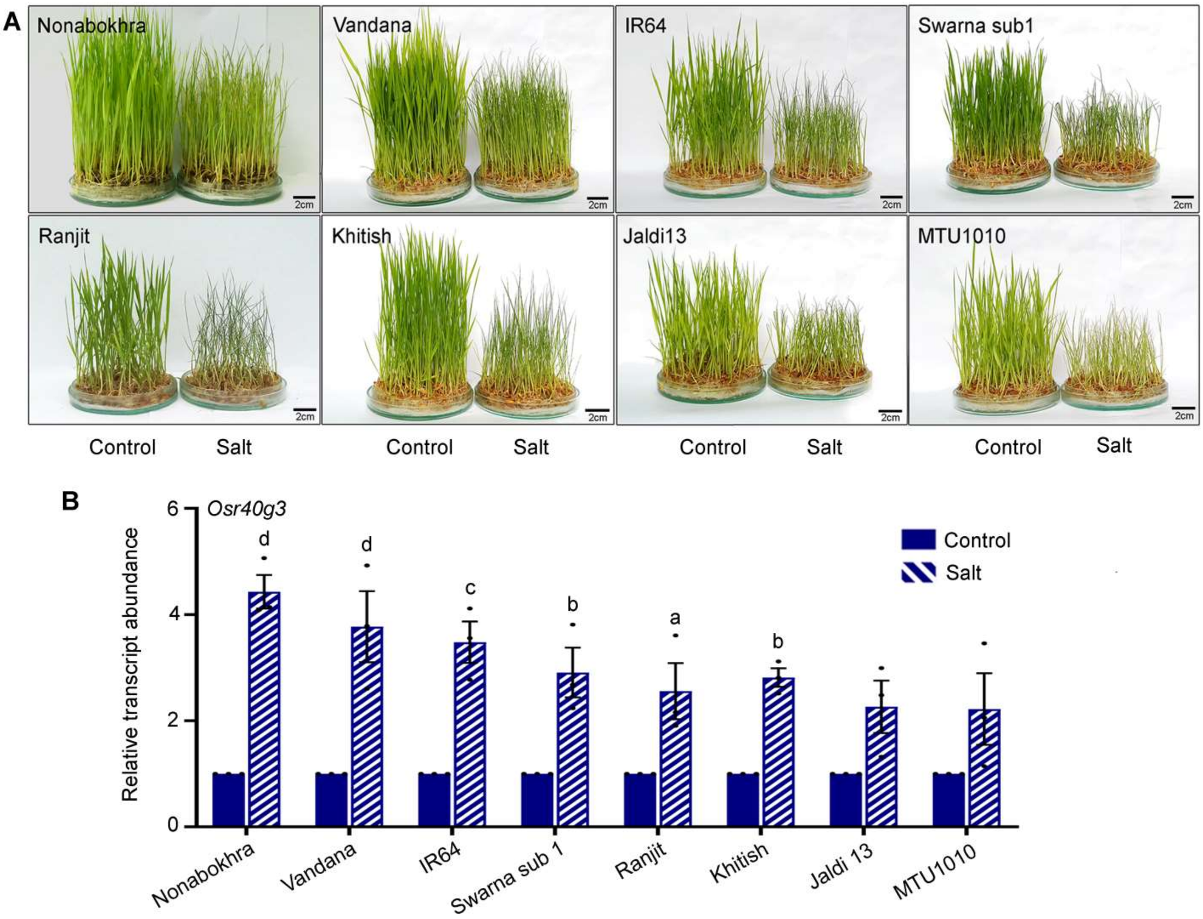


**Figure S1**. **Salt stress assay of 8 *indica* rice cultivars**. 15-days-old seedlings were treated with 200 mM NaCl solution for 5 d and morphological changes were analyzed. (A) Morphology of 8 *indica* rice cultivars in response to salt stress. (B) After salt treatment, samples were used for qRT-PCR analysis to study the relative transcript abundance of *Osr40g3* gene. The experiment was independently repeated thrice and results were represented as mean ± SEM. Statistical difference between samples under control and salt stress was denoted by different letters at $P<0.05$ (a), $P<0.01$ (b), $P<0.001$ (c) and $P<0.0001$ (d).

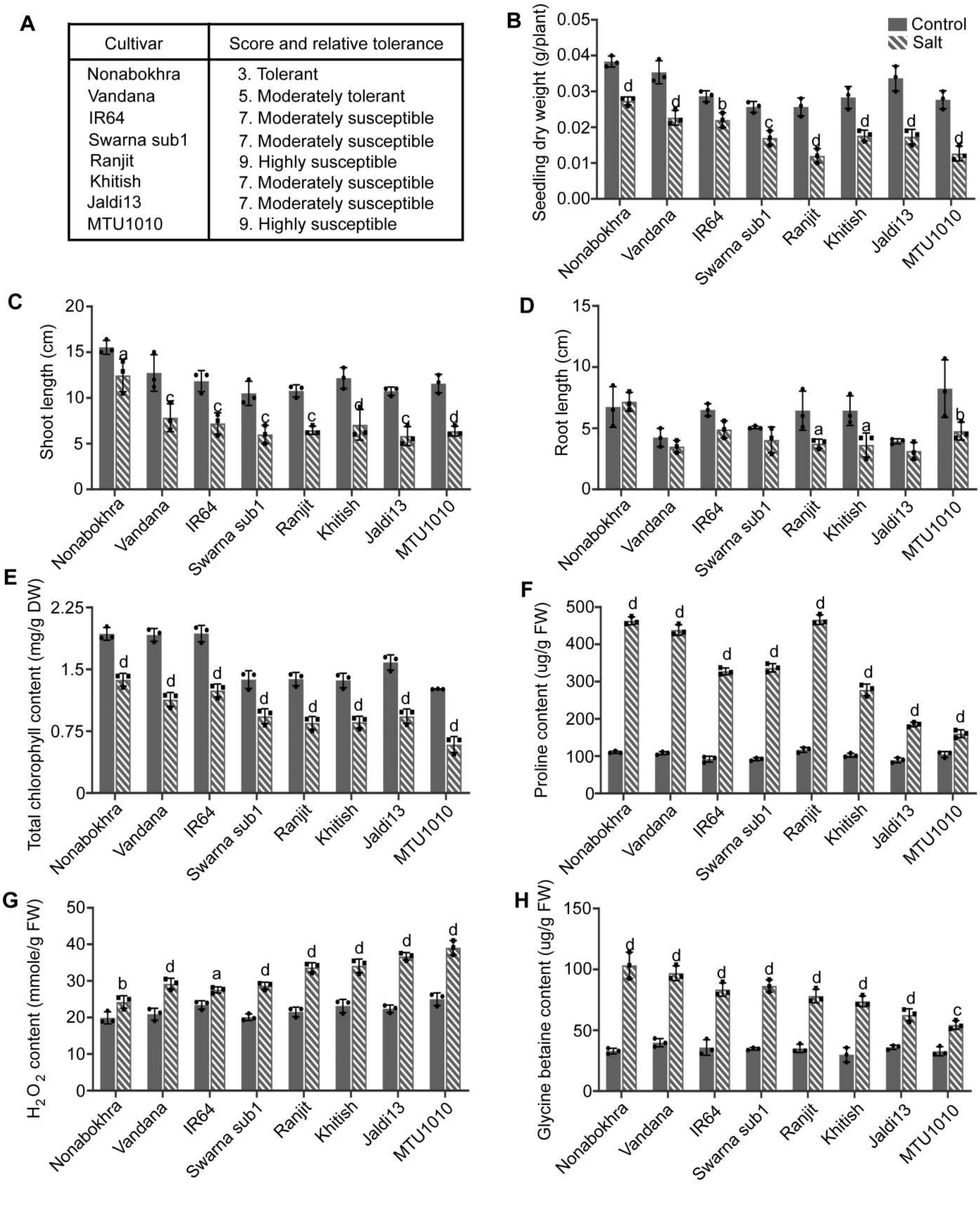


**Figure S2. Morphological and biochemical analyses of the 8 *indica* rice cultivars in response to salt stress.** (A) Scoring of salt stress response according of IRRI (2013), (B) seedling dry weight, (C) shoot length, (D) root length, (E) chlorophyll content, (F) proline content, (G) $H_2O_2$ content, and (H) glycine betaine content. Three biological replicates were considered for each experiment and results were represented as mean ± SEM. Statistical difference between samples under control and salt stress was denoted by different letters at P<0.05 (a), P<0.01 (b), P<0.001 (c) and P<0.0001 (d).

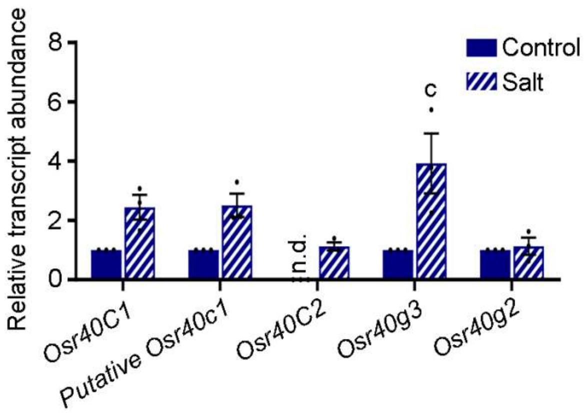


**Figure S3. Relative transcript abundance of 5 different *Os*R40 family genes from rice shoots in response to salt stress condition.** *Osr40g3* displayed highest induction in response to salt stress. The experiment was independently repeated thrice and results were represented as mean ± SEM. Statistical difference under control and salt stress was denoted by different letters at P<0.001 (c). n.d.: not detected.

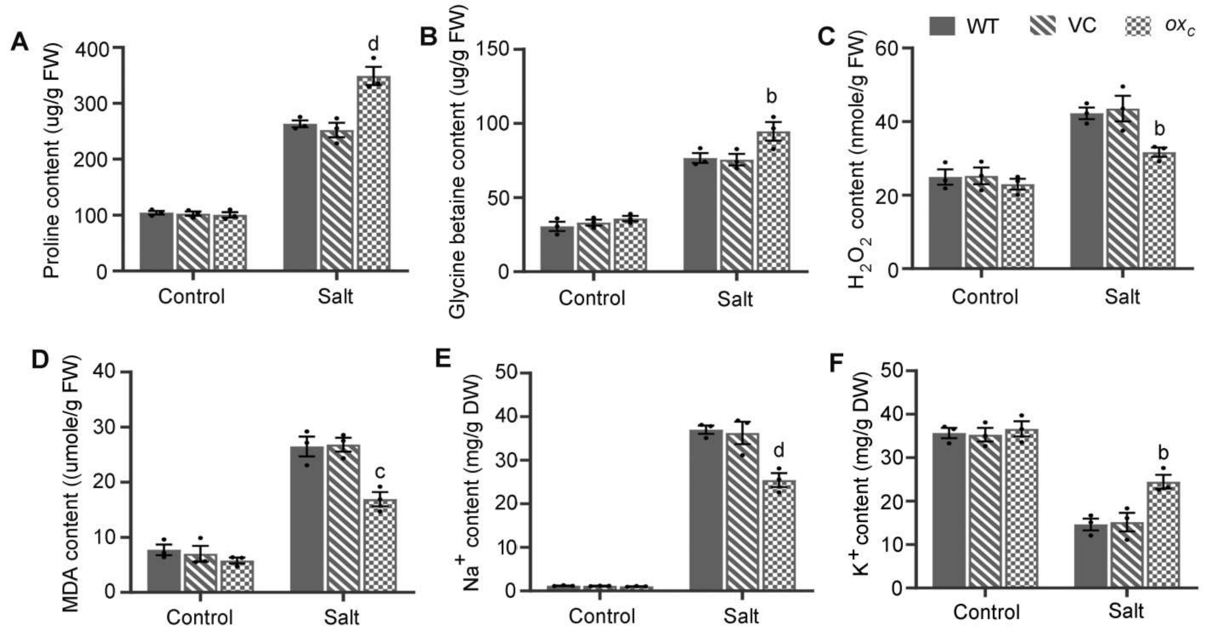


**Figure S4. Biochemical analyses of the transgenic rice lines constitutively overexpressing the *Osr40g3* gene (*ox*$_c$ lines) in response to salt stress.** (A) Proline content, (B) glycine betaine content, (C) $H_2O_2$ content, (D) MDA content, (E) sodium ($Na^+$) content, and (F) potassium ($K^+$) content. The experiment was performed with 3 independent transgenic lines ($T_0$) and results were represented as mean ± SEM. Statistical difference between the WT, VC, and *ox*$_c$ lines was denoted by different letters at P<0.01 (b), P<0.001 (c) and P<0.0001 (d).

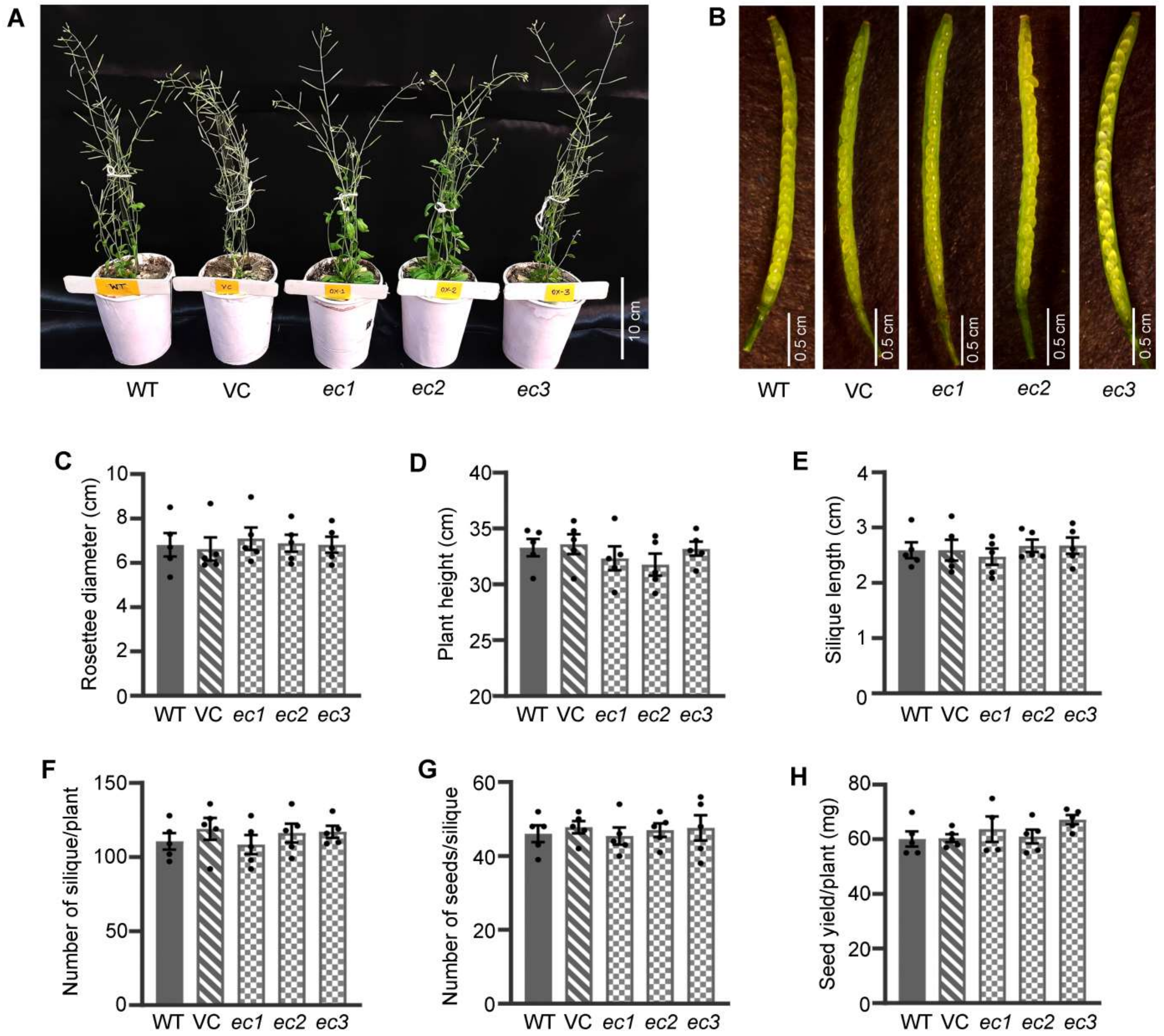


**Figure S5**. **Morphological characterization of transgenic *Arabidopsis* lines ectopically expressing *Osr40g3***. Different morphological parameters of 3 independent transgenic lines harboring the *35S::Osr40g3* construct (*ec* lines) were recorded and compared with WT and VC (*35S::GFP*) plants. The transgenic lines exhibited no phenotypic abnormalities. (A) WT, VC and *ec* lines at reproductive stage, (B) siliques, (C) rosette diameter, (D) plant height, (E) silique length, (F) number of siliques per plant, (G) number of seeds per silique, and (H) seed yield per plant. Five biological replicates were considered for each line ($T_2$) and results were represented as mean ± SEM. No statistically significant difference was observed between WT, VC and *ec* lines.

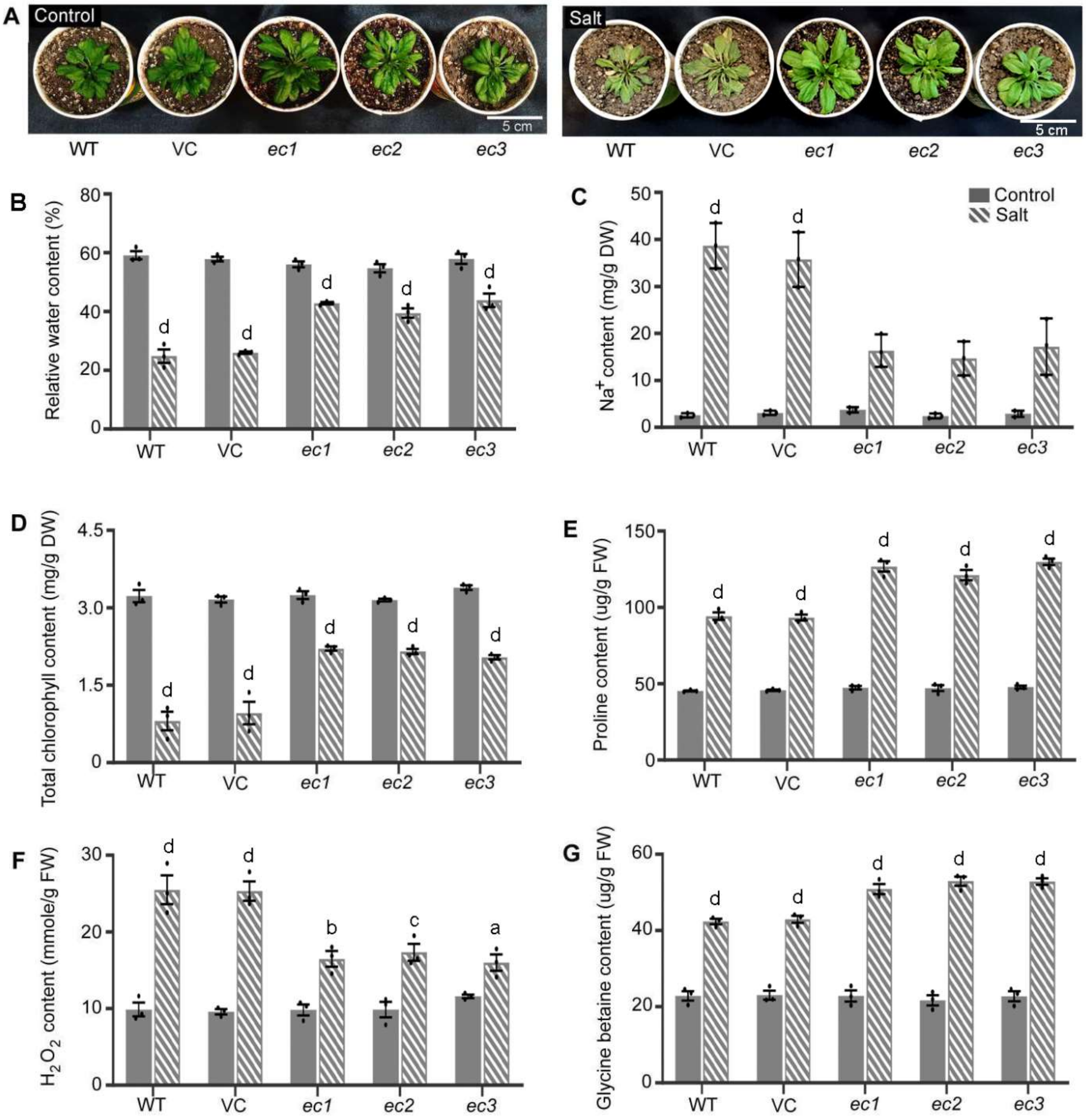


**Figure S6. Response of transgenic *Arabidopsis* lines ectopically expressing *Osr40g3* gene under salt stress condition.** The WT, VC (*35S::GFP*) and 3 independent transgenic lines harboring the *35S::Osr40g3* construct (*ec* lines) were exposed under salt stress for 5 d and analyzed. The transgenic lines exhibited improved salt stress tolerance over WT and VC lines. (A) Morphological response under salt stress, (B) relative water content, (C) sodium ($Na^+$) content, (D) chlorophyll content, (E) proline content, (F) $H_2O_2$ content, and (G) glycine betaine content. The experiment was independently repeated thrice with $T_2$ plants and results were represented as mean ± SEM. Statistical difference between the samples under control and salt stress was denoted by different letters at $P<0.05$ (a), $P<0.01$ (b), $P<0.001$ (c) and $P<0.0001$ (d).

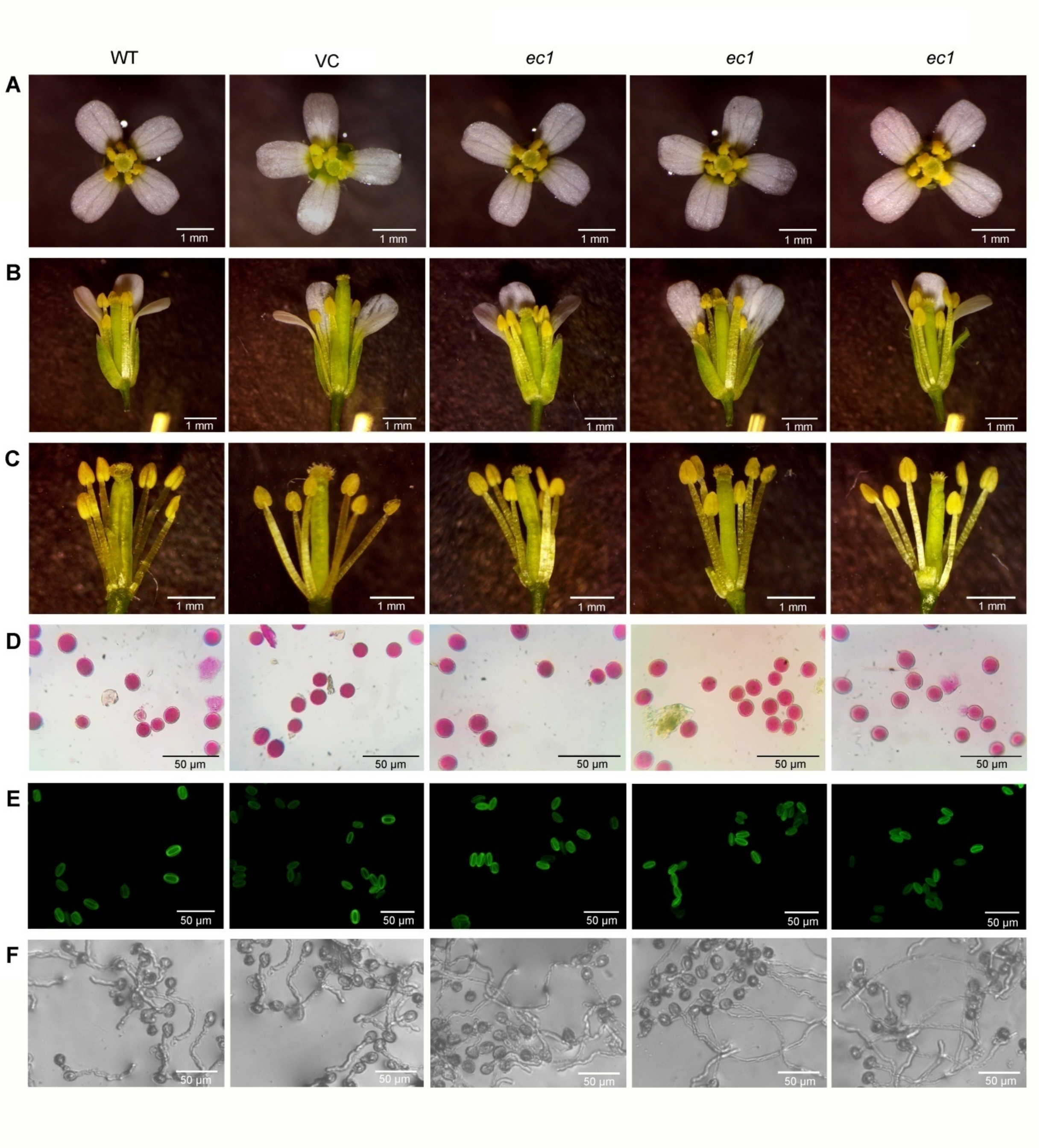


**Figure S7**. **Floral morphology of transgenic *Arabidopsis* lines ectopically expressing *Osr40g3*.** The floral morphology of 3 independent transgenic lines harboring the *35S::Osr40g3* construct (*ec* lines) were recorded and compared with WT and VC (*35S::GFP*) plants. No morphological alteration was observed between the transgenic, VC and WT flowers. (A) Entire flower, (B) dissected flower, (C) stamens with pistil, (D) pollen viability assay by alexander staining method, (E) pollen viability assay by FDA staining method, and (F) *in vitro* pollen germination assay. Five biological replicates were considered for each line ($T_2$). For pollen viability and *in vitro* germination assay, at least 500 pollen grains from each line were studied.

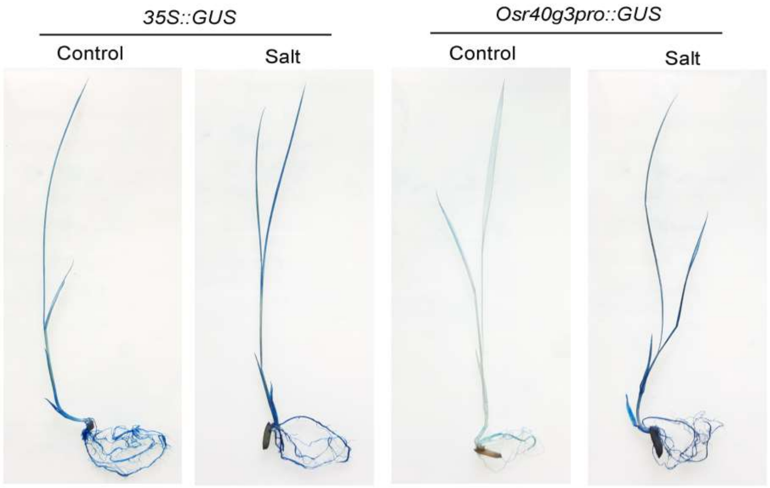


**Figure S8**. **Histochemical GUS assay for *Osr40g3* promoter activity in response to salt stress**. The VC line harboring the *35S::GUS* construct was used as a control. The experiment was independently repeated thrice.

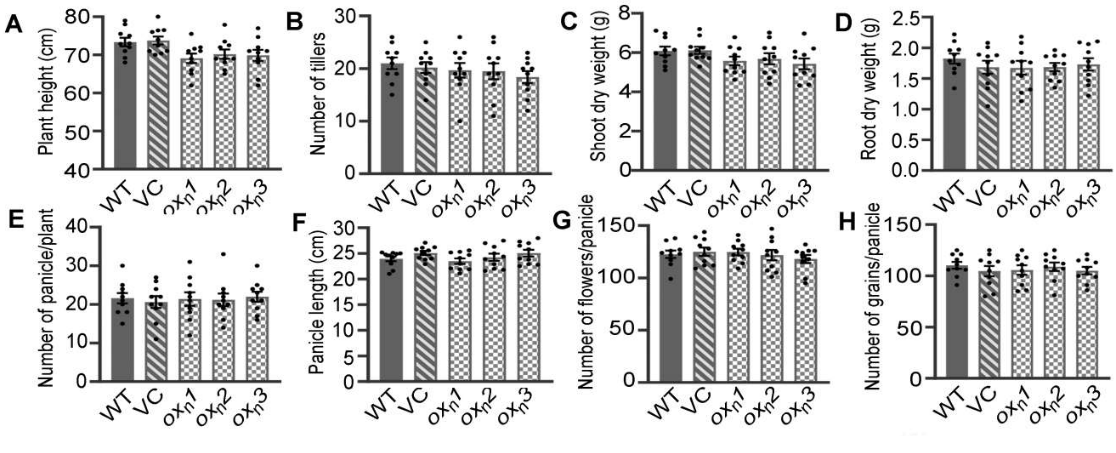


**Figure S9. The transgenic rice plants overexpressing *Osr40g3* under control its native promoter (*$ox_n$* lines) displayed no phenotypic abnormality**. Different morphological parameters of 3 independent transgenic lines (*$ox_n$*) harboring the *Osr40g3pro::Osr40g3* construct were recorded and compared with WT and VC (*35S::GFP*) plants. (A) plant height, (B) number of tillers, (C) shoot dry weight, (D) root dry weight, (E) numbers of panicles/plant, (F) panicle length, (G) numbers of flowers/panicle, and (H) number of grains/panicle. For morphological analysis, 10 biological replicates were considered for each line ($T_1$). Results were represented as mean ± SEM.

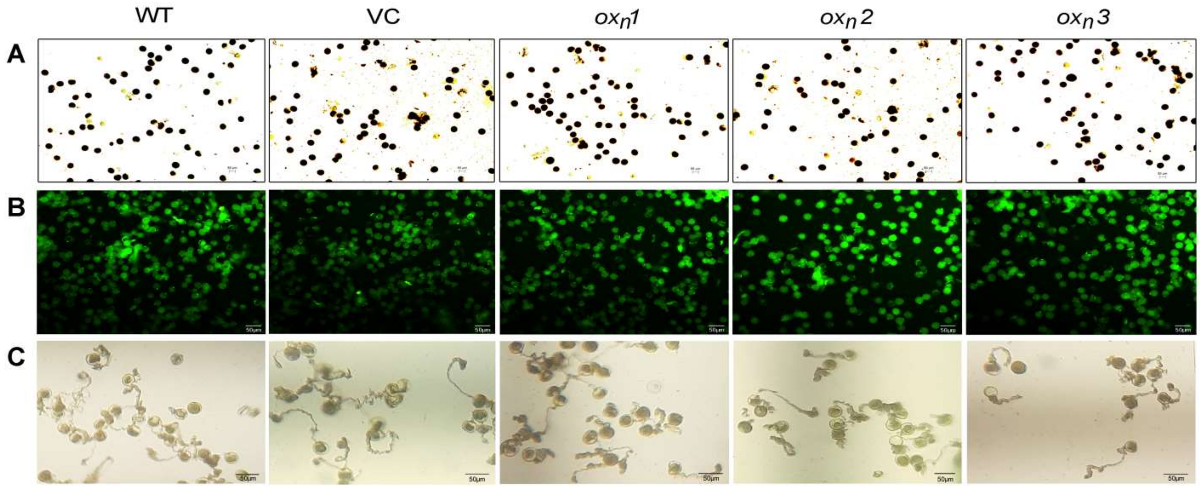


**Figure S10**. **Pollen viability of transgenic rice lines overexpressing *Osr40g3* under control of native promoter.** The pollen viability of 3 independent transgenic lines harboring the *Osr40g3pro::Osr40g3* construct (*ox*$_n$ lines) along with the WT and VC (*35S::GFP*) plants were analyzed. No significant difference was observed in the pollen viability and germination efficiency of the transgenic pollen grains as compared with the WT and VC lines. (A) Pollen viability by $I_2$-KI staining method, (B) pollen viability by FDA staining method, and (C) *in vitro* pollen germination assay. At least 500 pollen grains from each line ($T_1$) were studied.

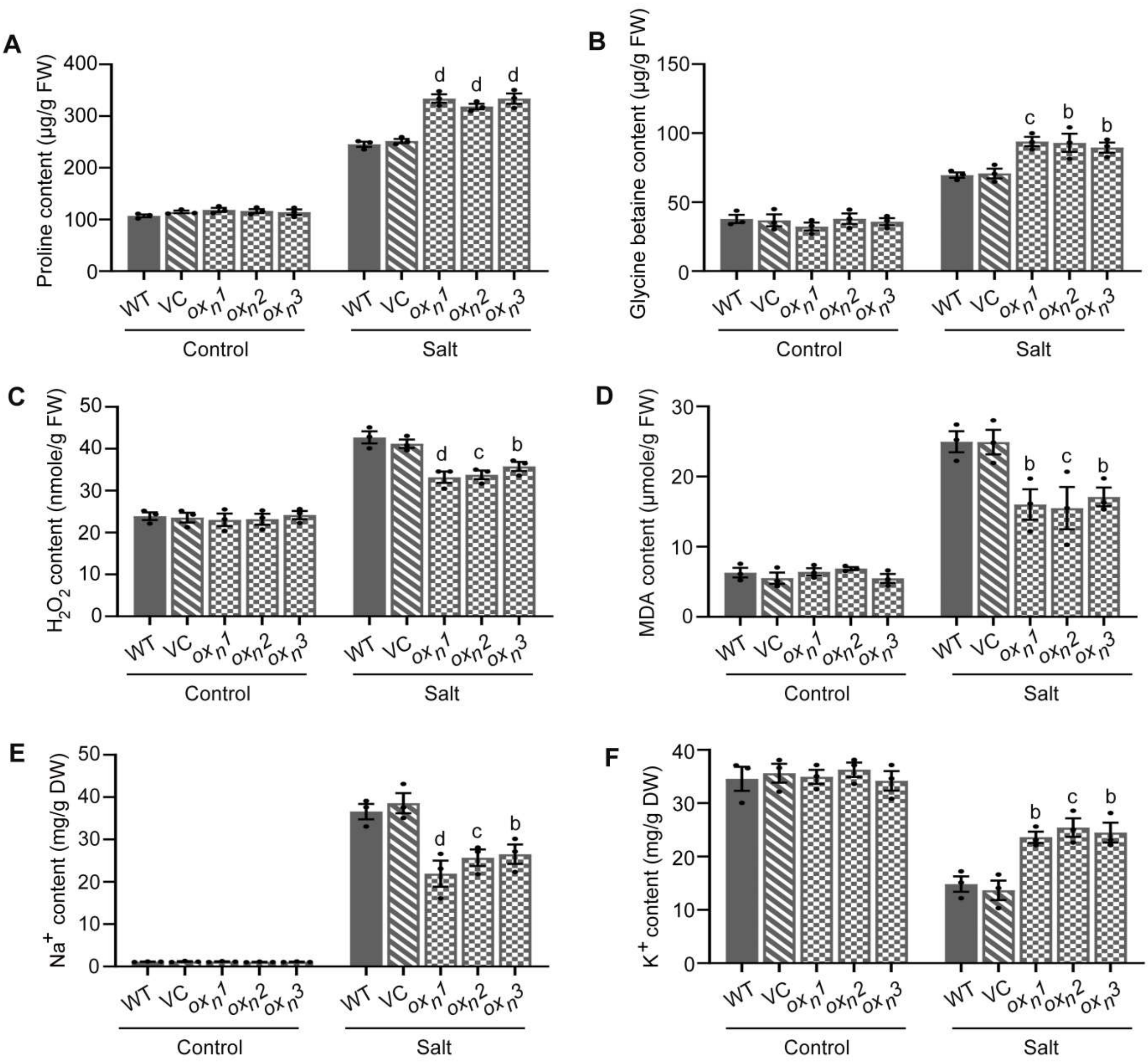


**Figure S11. Biochemical analyses of the transgenic rice lines overexpressing the *Osr40g3* gene (*$ox_n$* lines) under control of the native promoter in response to salt stress.** (A) Proline content, (B) glycine betaine content, (C) $H_2O_2$ content, (D) MDA content, (E) sodium ($Na^+$) content, and (F) potassium ($K^+$) content. The experiment was independently repeated thrice with 3 independent transgenic lines ($T_1$) and results were represented as mean ± SEM. Statistical difference between the WT, VC, and transgenic lines was denoted by different letters at P<0.01 (b), P<0.001 (c) and P<0.0001 (d).

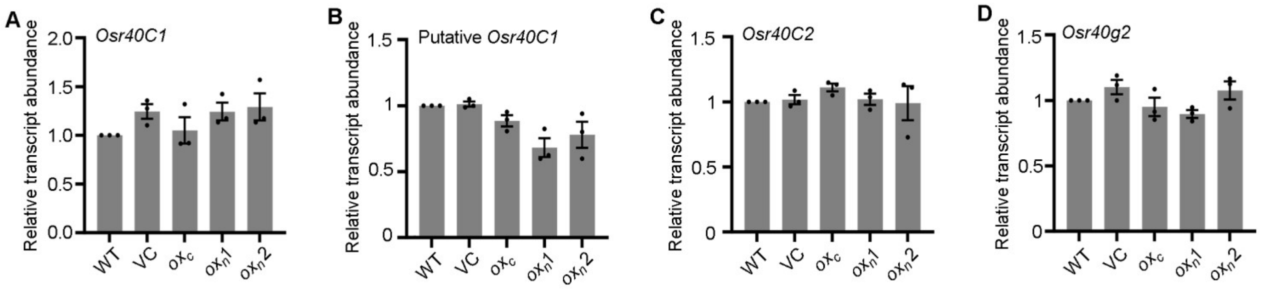


**Figure S12. Relative transcript abundance of *OsR40* genes in WT, VC, *$ox_c$* and *$ox_n$* lines plant.** The experiment was independently repeated thrice and results were represented as mean ± SEM.

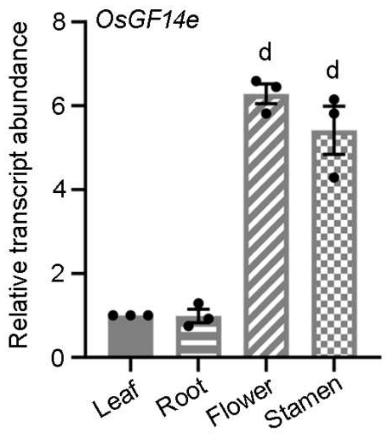


**Figure S13. Relative transcript abundance of *OsGF14e* in leaf, root, flower and stamen of WT plant.** The experiment was independently repeated thrice and results were represented as mean ± SEM. Statistical difference of the samples was denoted at P<0.0001 (d).

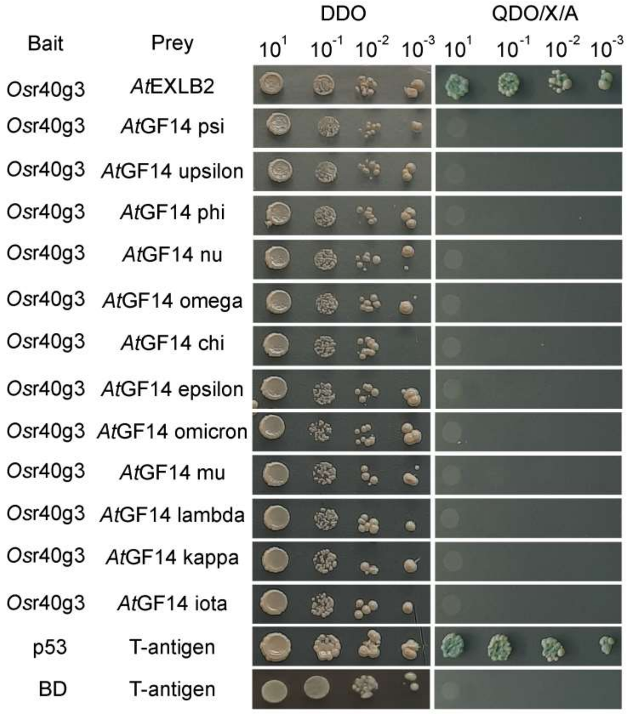


**Figure S14. Yeast two-hybrid assay for the interaction of *Os*r40g3 with the *Arabidopsis* orthologs of *Os*GF14e and *Os*EG45 proteins.** *Os*r40g3 interacted with *At*EXLB2, the orthologs of *Os*Eg45, while no interaction between *Os*r40g3 and the orthologs of *Os*GF14e was observed. The interaction of p53 protein with T-antigen was used as a positive control. The interaction of the empty BD with T-antigen was used as a negative control.

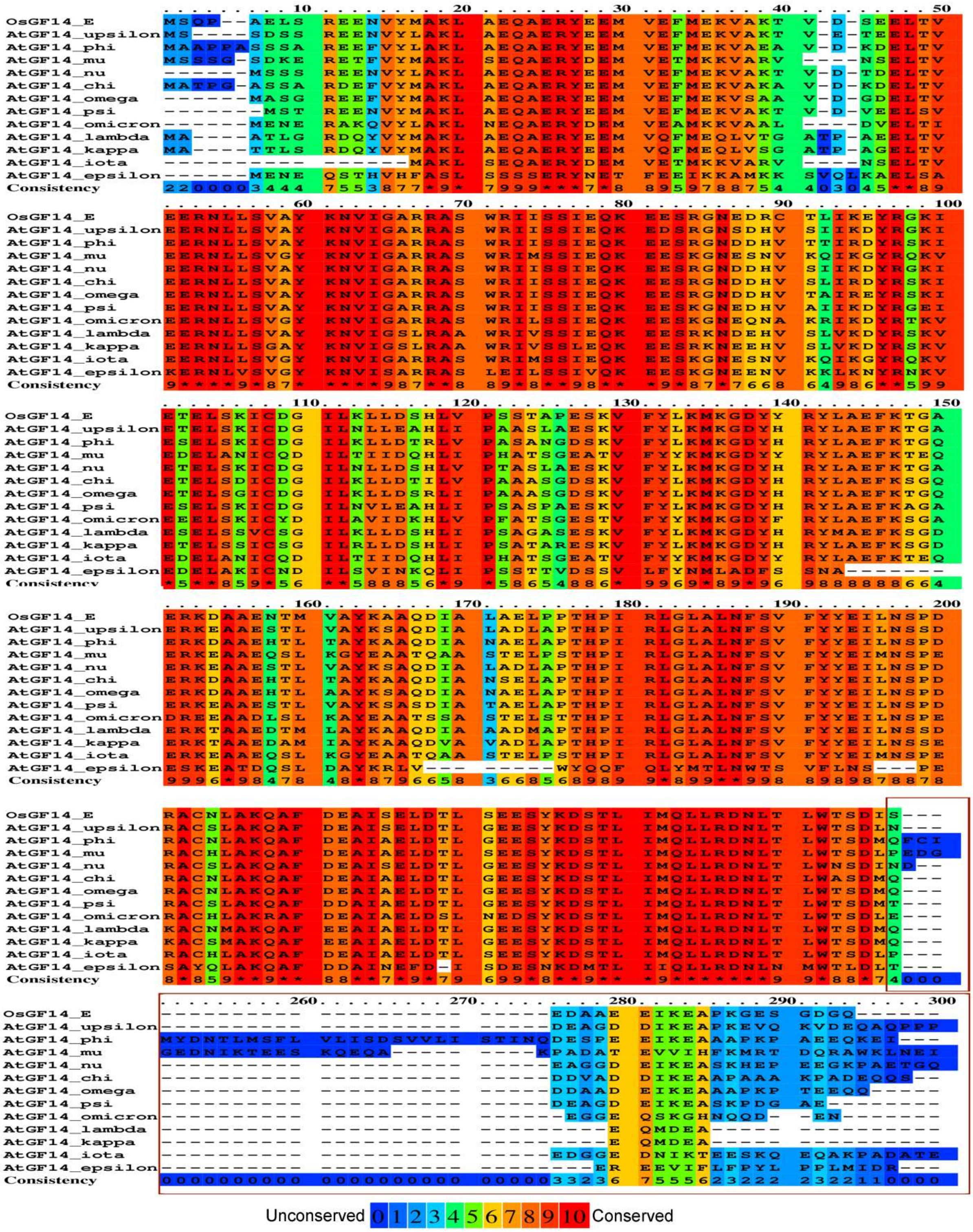

**Figure S15. Multiple sequence alignment of *Os*GF14e protein with *Arabidopsis* GF14 family members showing structural variability at the C-terminal end.**

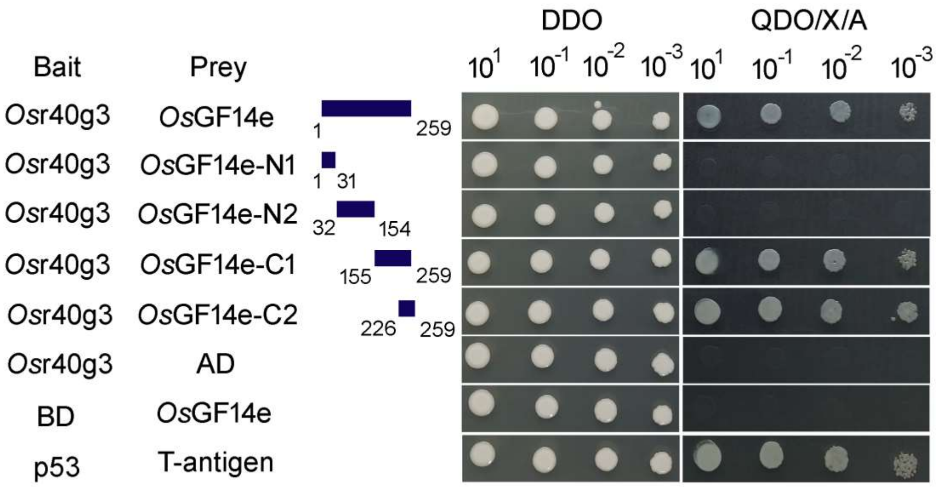


**Figure S16. Yeast two-hybrid assay for the interaction of *Os*r40g3 with N-terminal and C-terminal domains of *Os*GF14e.** Positive interaction was observed only in case of C-terminal domain of *Os*GF14e. The interaction of p53 protein with T-antigen was used as a positive control. The interaction of the empty BD with *Os*GF14e as well as *Os*r40g3 with empty AD was used as a negative control.

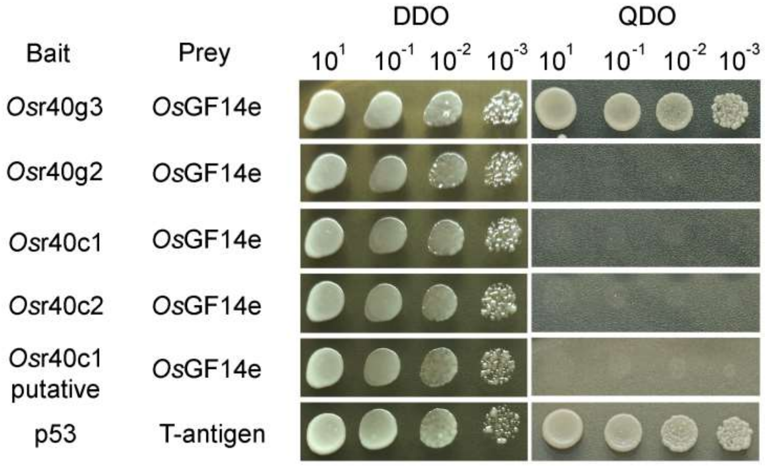


**Figure S17. Yeast two-hybrid assay for the interaction of *Os*GF14e with all the *Os*r40 family proteins.** Positive interaction was observed only in case of *Os*r40g3. The interaction of p53 protein with T-antigen was used as a positive control.

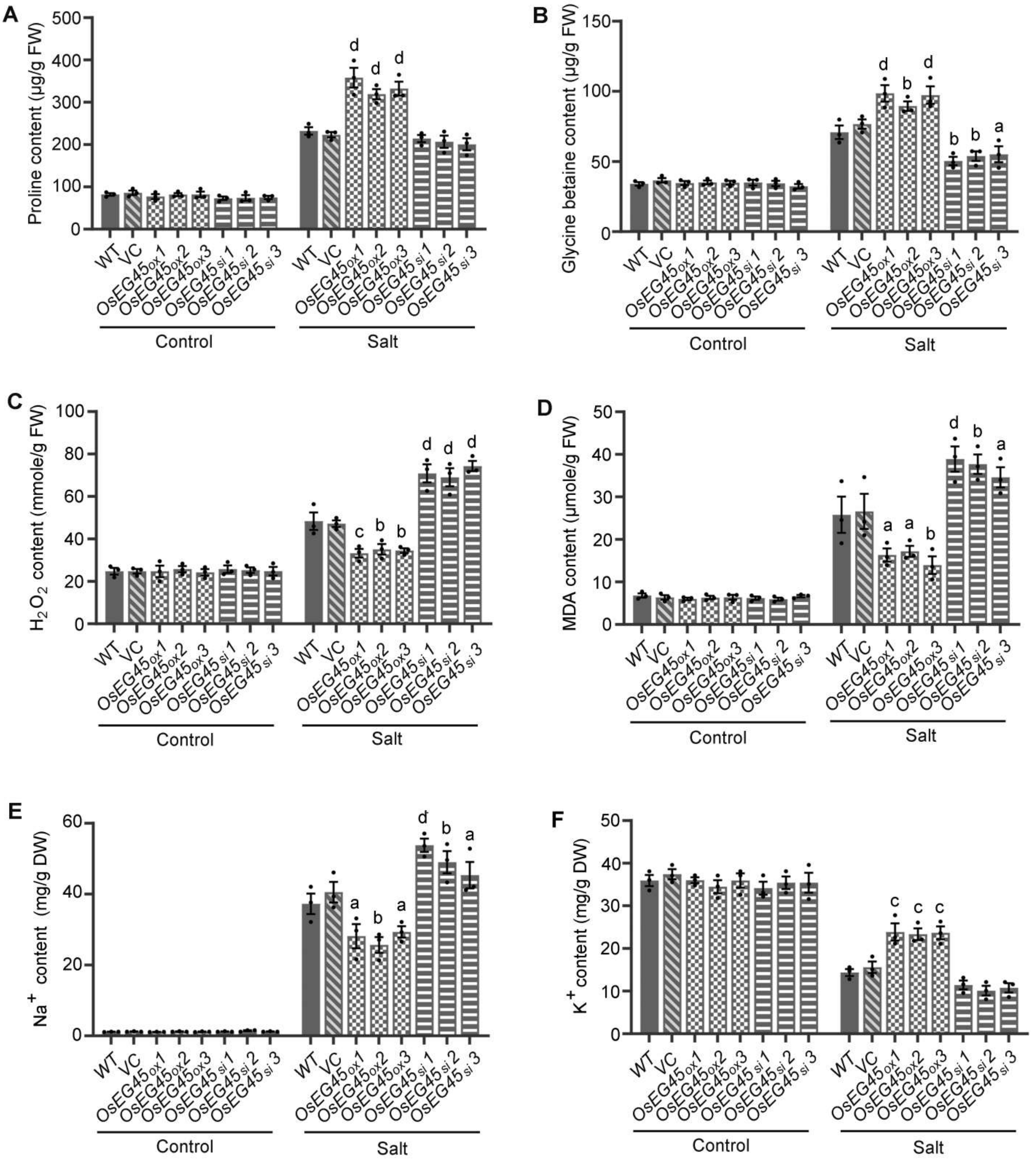


**Figure S18. Biochemical analyses of the *OsEG45* overexpressing (*OsEG45$_{ox}$* lines) and silencing (*OsEG45$_{si}$* lines) transgenic rice lines in response to salt stress.** (A) Proline content, (B) glycine betaine content, (C) $H_2O_2$ content, (D) MDA content, (E) sodium ($Na^+$) content, and (F) potassium ($K^+$) content. The experiment was independently repeated thrice with three independent transgenic lines ($T_1$) and results were represented as mean ± SEM. Statistical difference between the WT, VC, and transgenic lines was denoted by different letters at P<0.05 (a), P<0.01 (b), P<0.001 (c) and P<0.0001 (d).

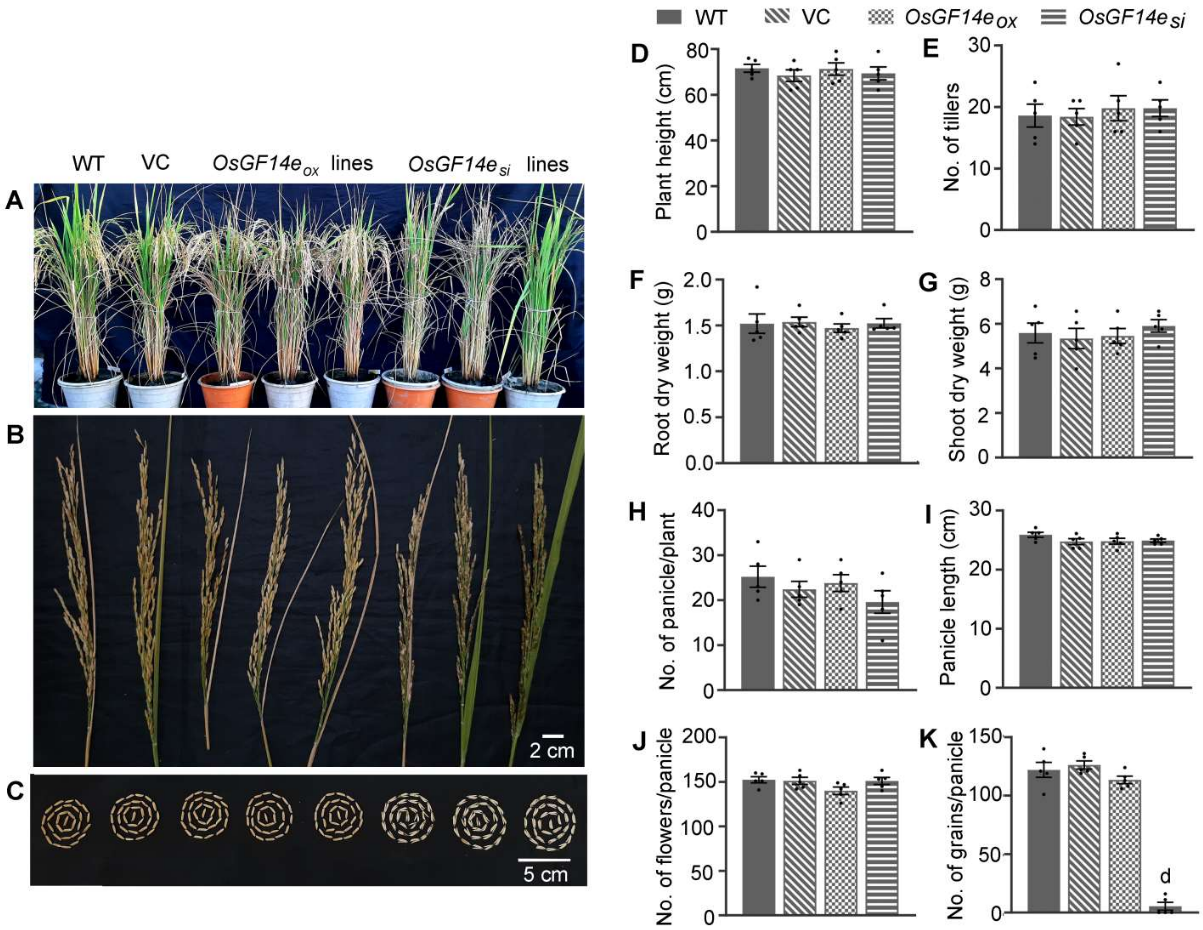


**Figure S19**. **Morphological characterization of the *OsGF14e* overexpressing (*OsGF14e$_{ox}$* lines) and silencing (*OsGF14e$_{si}$* lines) transgenic rice lines**. Different morphological parameters of 5 independent *OsGF14e* overexpressing (*OsGF14e$_{ox}$* lines) and silencing (*OsGF14e$_{si}$* lines) transgenic rice lines were recorded and compared with WT and VC (*35S::GFP*) plants. The *OsGF14e$_{ox}$* lines exhibited no phenotypic abnormalities while the *OsGF14e$_{si}$* lines failed to set seeds. (A) WT, VC, *OsGF14e$_{ox}$* and *OsGF14e$_{si}$* lines at reproductive stage, (B) panicles, (C) seed morphology, (D) plant height, (E) number of tillers, (F) root dry weight, (G) shoot dry weight, (H) number of panicles per plant, (I) panicle length, (J) number of flowers per panicles, and (H) number of seeds per panicle. Five independent transgenic lines ($T_0$) were considered for each case and results were represented as mean ± SEM. Statistical difference between the WT, VC and transgenic lines was denoted by P<0.0001 (d).

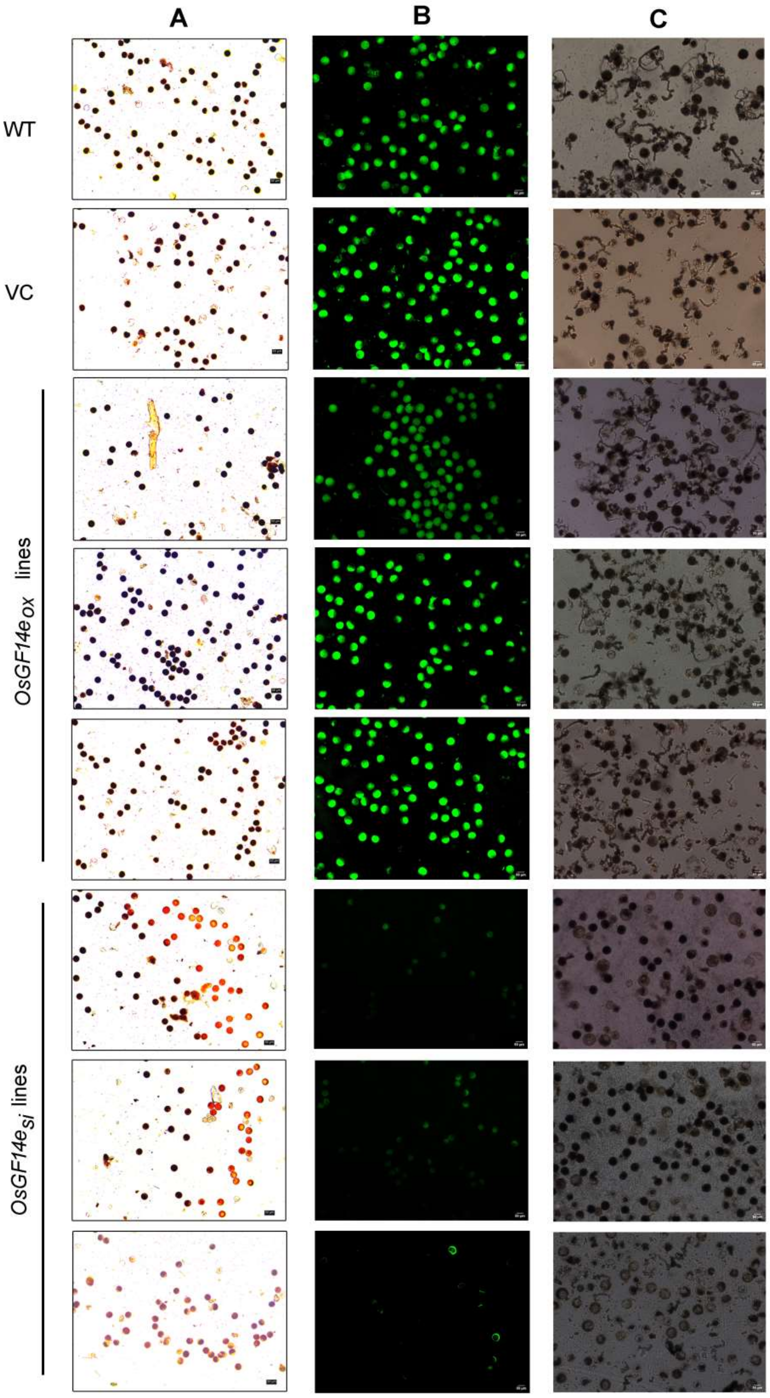


**Figure S20**. **Pollen viability of the *OsGF14e* overexpressing (*OsGF14e$_{ox}$* lines) and silencing (*OsGF14e$_{si}$* lines) transgenic rice lines.** The pollen viability of 3 independent *OsGF14e* overexpressing (*OsGF14e$_{ox}$* lines) and silencing (*OsGF14e$_{si}$* lines) lines were analyzed. No significant difference was observed in the pollen viability and germination efficiency of the *OsGF14e*$_{ox}$ lines as compared with the WT and VC lines while the *OsGF14e*$_{si}$ lines displayed severe pollen sterility. (A) Pollen viability by $I_2$-KI staining method, (B) pollen viability by FDA staining method, and (C) *in vitro* pollen germination assay. At least 500 pollen grains from each line ($T_0$) were studied.

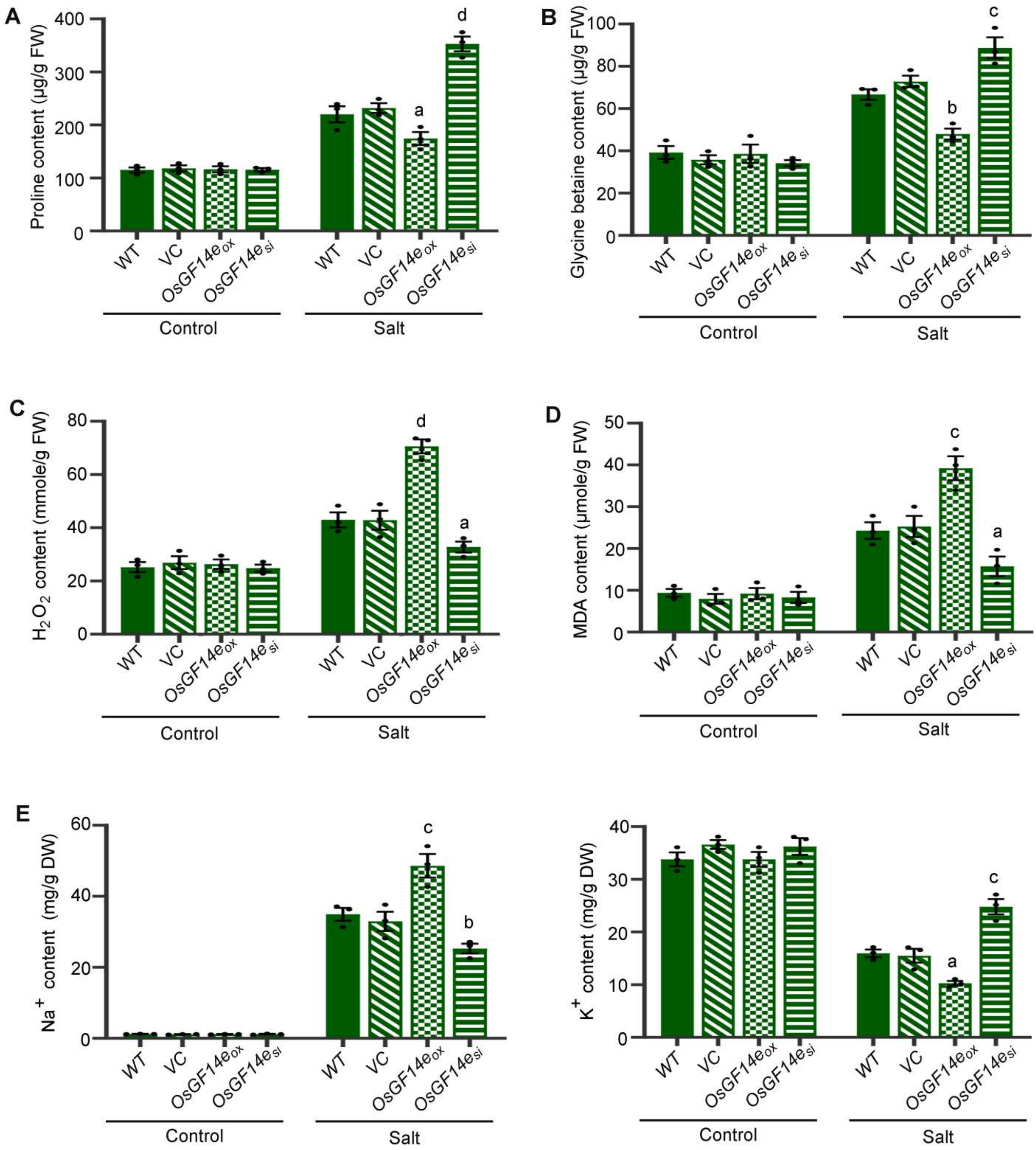


**Figure S21. Biochemical analyses of the *OsGF14e* overexpressing (*OsGF14e*$_{ox}$ lines) and silencing (*OsGF14e*$_{si}$ lines) transgenic rice lines in response to salt stress.** (A) Proline content, (B) glycine betaine content, (C) $H_2O_2$ content, (D) MDA content, (E) sodium ($Na^+$) content, and (F) potassium ($K^+$) content. Three independent transgenic lines ($T_0$) were considered in each case and results were represented as mean ± SEM. Statistical difference between the WT, VC, and transgenic lines was denoted by different letters at P<0.01 (b), P<0.001 (c) and P<0.0001 (d).

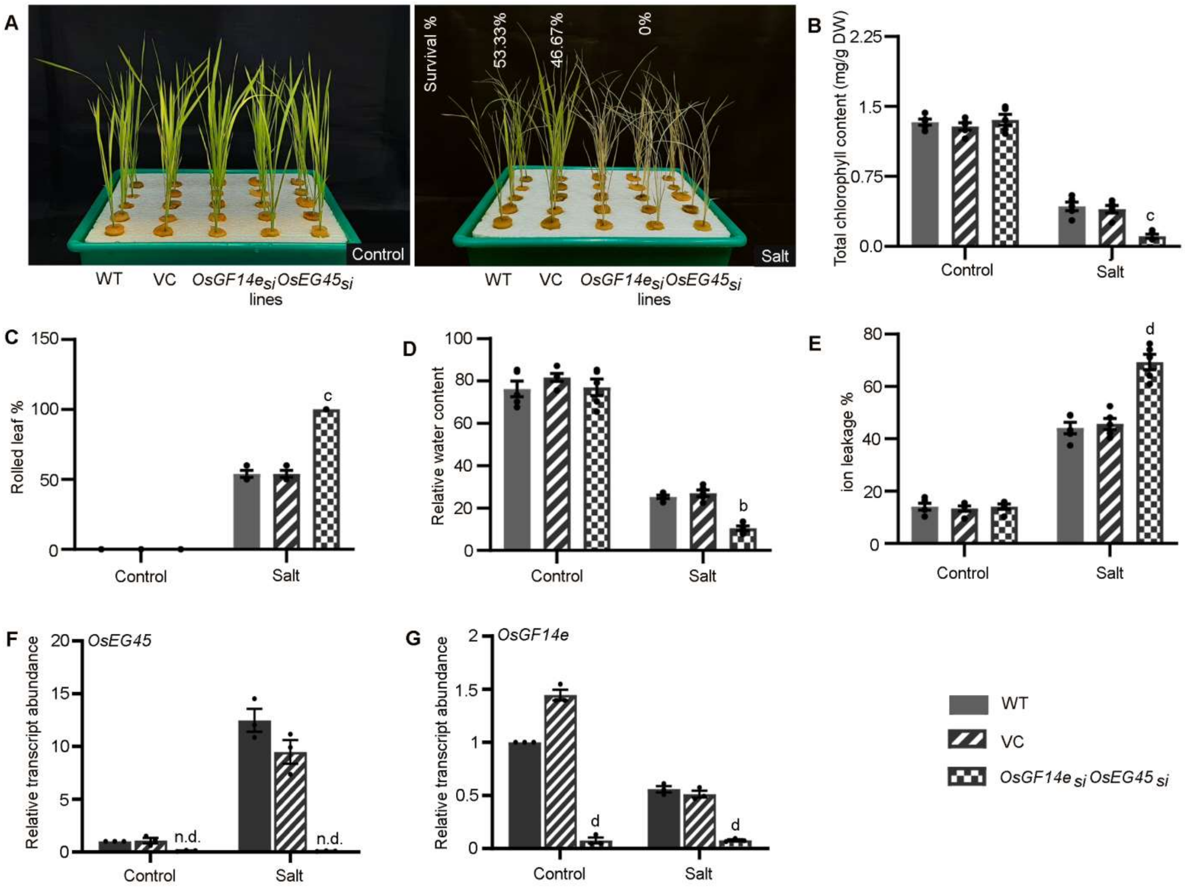


**Figure S22. Salt stress assay of the *$OsGF14e_{si}OsEG45_{si}$* double mutant.** 30-days-old hydroponically grown WT, VC and independent transgenic lines of the *$OsGF14e_{si}OsEG45_{si}$* double mutant were exposed to salt stress for 5 d and analyzed. (A) Morphological response under salt stress, (B) total chlorophyll content, (C) rolled leaf percentage, (D) relative water content, (E) relative ion leakage percentage, and relative transcript abundance of (F) *OsEG45* and (G) *OsGF14e* genes in response to salt stress. The stress assay performed with 15 independent lines ($T_0$) and subsequent analyses were performed with 5 independent lines ($T_0$). The relative transcript abundance of *OsEG45* and *OsGF14e* genes was analyzed in 3 independent *$OsGF14_{si}OsEG45_{si}$* lines. Results were represented as mean ± SEM and the statistical difference between WT, VC and transgenic lines was denoted by different letters at P<0.01 (b), P<0.001 (c), and P<0.0001 (d). n.d.: not detected.

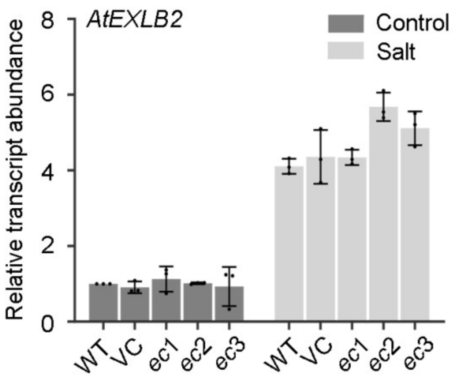


**Figure S23. Relative transcript abundance of *AtEXLB2* in WT, VC, and *ec* lines.** The experiment was independently repeated thrice and results were represented as mean ± SEM.

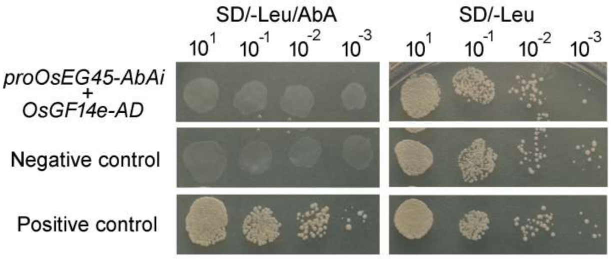


**Figure S24. Yeast one-hybrid assay for the interaction of *Os*GF14e with the *OsEG45* promoter.** No interaction was observed. The interaction of p53 protein with pAbAi was used as a positive control. The interaction of *Os*GF14e with pAbAi was used as a negative control.

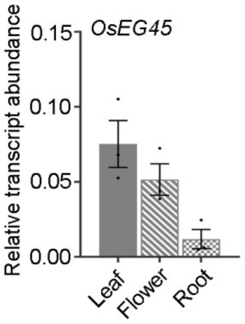


**Figure S25. Relative transcript abundance of *OsEG45* in different tissues of WT rice.** The experiment was independently repeated thrice and results were represented as mean ± SEM.

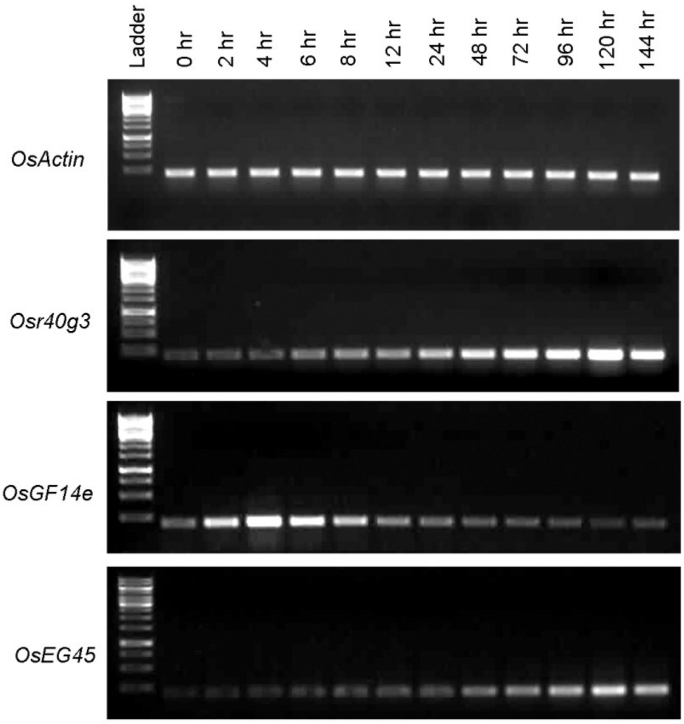


**Figure S26. Expression analysis of *Osr40g3*, *OsGF14e*, and *OsEG45* genes from leaves in response to salt stress on a temporal landscape.** The semi-qRT-PCR analysis was independently repeated thrice.
