## Supplementary Table for "*Os*r40g3 imparts salt tolerance by regulating GF14e-mediated gibberellin metabolism to activate EG45 in rice"

**Supplemental Table S1.** Estimation of soil salt concentration before and after salinity stress

| **Sample** | **Sodium (mg/kg)** | **Potassium (mg/kg)** | **Chloride (mg/kg)** | **EC (dS/m)** |
|---|---|---|---|---|
| Soil before salt treatment | 992.43 ± 11 | 3978.75 ± 77 | 309.84 ± 6 | 1.969±0.115 |
| Soil after salt treatment | 2366.67 ± 36 | 3323.18 ± 23 | 3357.50 ± 69 | 9.638±0.402 |

**Supplemental Table S2.** Primers used in this study

| Gene name | Loci number | Primers |
|---|---|---|
| *Osr40g3* | *LOC112936024* | Forward: 5'GAATTCATGGACTTTTACGGGCGGC3' |
| | | Reverse: 5'GGATCCTCAGTAGTAGGGCTGGATCT3' |
| *Osr40g3 (partial)* | *LOC112936024* | Forward: 5'AGCAGTACGGCGGGTATG3' |
| | | Reverse: 5'GTCCTTGATCCAATGCTGGT3' |
| *Osr40g3 promoter* | *LOC112936024* | Forward: 5'GGATCCAGTTGCCATGCATCCCATTGA3' |
| | | Reverse: 5'AGATCTACTCGCTAGCTACCTAGCTAA3' |
| *OsActin1* | *LOC4333919* | Forward- 5'CCGTCCTCCTGCTTGTTTCT3' |
| | | Reverse: 5'TGGTACCCTCATCAGGCATC3' |
| *AtActin* | *At3G18780* | Forward- 5'GCACCCTGTTCTTCTTACCG3' |
| | | Reverse: 5'AACCCTCGTAGATTGGCACA 3' |
| *Osr40c2* | *LOC_Os07g48460.1* | Forward: 5'TGAAGCTGGTGCCGTACA3' |
| | | Reverse: 5'GTATGCCTGGTGCCCGTAT3' |
| *Osr40g2* | *LOC4344319* | Forward: 5'CAAGGGCGACAACCAGAG3' |
| | | Reverse: 5'CTTCCTCGTCCCTGATCTTG3' |

| | | |
|---|---|---|
| *Osr40c1* | *LOC4332715* | Forward: 5’CCATCGTGCTCTGGGAGT3’ |
| | | Reverse: 5’TCCTCGTCCTTGATGCTGTT3’ |
| *Osr40c1putative* | *LOC_Os01g01450.2* | Forward: 5’CCTACGGCCACGACTCAG3’ |
| | | Reverse: 5’GGCCTTGCAGTAGATCCTGT3’ |
| *Bar* | *AF218816_2* | Forward: 5’AAGCACGGTCAACTTCCGTA3’ |
| | | Reverse: 5’GAAGTCCAGCTGCCAGAAAC3’ |
| *OsGF14e* | *LOC4329775* | Forward: 5’ACTAGT ATGTCGCAGCCTGCTGAG3’ |
| | | Reverse: 5’GGATCCTCACTGTCCATCTCCTGATTC3’ |
| *AtGF14chi* | *At4g09000* | Forward: 5’ATG GCG ACA CCA GGA GCT T3’ |
| | | Reverse: 5’TAA GGATTGTTGCTCGTCAGC3’ |
| *AtGF14omega* | *At1g78300* | Forward:5’ATG GCGTCTGGGCGT GAAG3’ |
| | | Reverse: 5’TCACTGCTGTTCCTCGGTC3’ |
| *AtGF14psi* | *At5g38480* | Forward:5’ATGTCGACAAGGGAAGAGAAT3’ |
| | | Reverse: 5’TTACTCGGCACCATCGGGC3’ |
| *AtGF14phi* | *At1g35160* | Forward: 5’ATGGCGGCACCACCAGCAT3’ |
| | | Reverse:5’TTAGATCTCCTTCTGTTCTTCAG 3’ |
| *AtGF14upsilon* | *At5g16050* | Forward:5’ATGTCTTCTGATTCGTCCCG3’ |
| | | Reverse: 5’TCACTGCGAAGGTGGTGGT3’ |
| *AtGF14lamda* | *At5g10450* | Forward:5’ATGGCGGCGACATTAGGCA3’ |
| | | Reverse: 5’TCAGGCCTCGTCCATCTGC3’ |
| *AtGF14nu* | *At3g02520* | Forward:5’ATGTCGTCTTCTCGGGAAGA3’ |
| | | Reverse: 5’TCACTGCCCTGTCTCAGCT3’ |
| *AtGF14kappa* | *At5g65430* | Forward: 5’ATGGCGACGACCTTAAGCAG3’ |
| | | Reverse: 5’TCAGGCCTCATCCATCTGCT3’ |
| *AtGF14mu* | *At2g42590* | Forward: 5’ATGGGTTCTGGAAAAGAGCG3’ |

| | | |
|---|---|---|
| | | Reverse: 5'ATTCGTCTTATGAGCATCGTCT3' |
| *AtGF14epsilon* | *At1g22300* | Forward:5'ATGGAGAATGAGAGGGAAAAG3' |
| | | Reverse: 5'TCATACCTCATCTTGAGGCTC3' |
| *AtGF14omicron* | *At1g34760* | Forward: 5'ATGGAGAACGAGAGAGCGAAGC3' |
| | | Reverse: 5' TTACTTACCTCCTTCCTCTAGATC3' |
| *AtGF14iota* | *At1g26480* | Forward: 5'ATGTCATCATCAGGATCCGACAA3' |
| | | Reversre: 5' TCAGTTCTCAGTGGCATCGGCA3' |
| *AtEXLB2* | *AT4G30380* | Forward:5'AGCTGCTCAAGGAAAAGCTG3' |
| | | Reverse:5'TCCCGCAGAAATCAACTAC3' |
| *OsEG45* | *LOC4347346* | Forward:5'TCT AGAATGACGACGATGGCTAAGGT3' |
| | | Reverse:5'GGATCCTTACACCTGTTGGTAGTCGAT3' |
| *OsEG45* promoter | *LOC4347346* | Forward:5'GAGCTC ACG TAG GCT CGA TGG TTA AAG A3' |
| | | Reverse:5' GGTACC CGC GAT CAA TTC GAT CGA TCA G3' |
| *OsGF14e*RNAi | *LOC4329775* | Forward:5'ACCA GGTCTC AGGAG CTG CTC CAG AGT CCA AGG TC3' |
| | | Reverse:5' ACCA GGTCTC ATCGT GCA GAA GCT GCA TGA TCA AA3' |
| *OsEG45*RNAi | *LOC4347346* | Forward:5'ACCA GGTCTC AGGAG CAC TTC GGT TGT CAT TGC AG3' |
| | | Reverse:5'ACCA GGTCTC ATCGT ATC TTC ACG GTG ACG CTC TG3' |
| *OsGf14e* c1ter | *LOC4329775* | Forward: 5' ATGGTGGCATACAAAGCCGC 3' |
| | | Reverse: 5' TCACTGTCCATCTCCTGATT 3' |
| *OsGf14e* c2 ter | *LOC4329775* | Forward: 5' ATGCAGCTTCTGCGTGATAA3' |
| | | Reverse: 5' TCACTGTCCATCTCCTGATT 3' |
| *OsGf14e* n1 ter | *LOC4329775* | Forward: 5' ATGTCGCAGCCTGCTGAGCT 3' |
| | | Reverse: 5' GGTGTTCTCAGCAGCATCCT 3' |
| *OsGf14e n2* ter | *LOC4329775* | Forward: 5' ATGTCGCAGCCTGCTGAGCT 3' |
| | | Reverse: 5' GATCAAAGTGCTGTCCTTGT 3' |
